# DEPP1 connects nutrient and oxygen availability to maintenance of muscle mass

**DOI:** 10.64898/2026.09.21.753276

**Authors:** Shariq Qayyum, Christopher Lavallee, Madhu Singh, Gregory A. Wyant

**Affiliations:** Massachusetts General Hospital, Cardiovascular Research Center, Division of Cardiology, Department of Medicine, Harvard Medical School, Charlestown, MA, 02129, USA; Broad Institute of MIT and Harvard, Cambridge, Massachusetts, USA

## Abstract

Nutrients and oxygen are sensed within the muscle to control growth and disruption of either signal is sufficient to lead to muscle atrophy^1^. While nutrient limitation is sensed via a conserved transcriptional atrophy program (commonly referred to as “atrogenes”) dictated via the Forkhead box O (FoxO) transcription factors, how low oxygen promotes muscle loss remains unknown^2^. Accordingly, the downstream mechanisms that initiate muscle loss when oxygen and nutrients are limiting are only partly understood^3^. Here, we find Hypoxia Inducible Factor (HIF), the master regulator of our adaptation to low oxygen, is necessary and sufficient to mediate muscle loss under hypoxia in mice. RNA sequencing in skeletal muscle isolated from starved or hypoxic mice identifies Decidual Protein Induced by Progesterone 1 (Depp1), which is induced in skeletal muscle when nutrients or oxygen is limiting via FoxO1 and HIF activation, respectively. Whole body Depp1 loss in mice reduces muscle loss under fasting and hypoxia and skeletal muscle Depp1 overexpression is sufficient to mediate muscle atrophy. Mechanistically, Depp1 localizes to the mitochondria and is necessary to control autophagy activation and mitochondrial degradation in skeletal muscle. Taken together, our studies nominate Depp1 as a new “atrogene” necessary for muscle loss under multiple atrophy scenarios involving FoxO and HIF.

**Significance:** Nutrients or oxygen are necessary to maintain muscle mass and when lost lead to muscle atrophy. While nutrient limitation has been shown to lead to muscle loss via the FoxO family of transcription factors, how we lose muscle mass when oxygen is limiting remains unknown. Here, we identify an overlapping atrophy mechanism that converges on Depp1, which is induced in skeletal muscle when nutrients or oxygen is limiting via the FoxO and Hypoxia Inducible Factor transcription factors. Mechanistically, Depp1 is necessary and sufficient to mediate muscle loss and is critical to maintain mitochondrial abundance under these conditions. These findings have the potential to inform how to preserve muscle mass under diverse pathophysiological scenarios where nutrients or oxygen are limiting.

## Introduction

Nutrients and oxygen are sensed within our muscle to control muscle growth^1^. Disruption of either signal occurs in multiple human pathological conditions, such as chronic starvation, heart failure, chronic bed rest, pulmonary arterial hypertension (PAH), and chronic obstructive pulmonary artery disease (COPD), and is sufficient to promote muscle atrophy^4–8^. Nutrient limitation (or starvation) leads to atrophy through activation of a common transcriptional program via the Forkhead box O (FoxO) transcription factors, comprised of Foxo1, 3, and 4, in the muscle^2^. How low oxygen (hypoxia) leads to muscle loss remains unknown. Accordingly, the downstream mechanisms that initiate muscle loss under nutrient and oxygen limitation are complex and still only partially understood.

Muscle atrophy involves the combined activation of two intracellular degradation pathways, the ubiquitin-proteasome system and autophagy, leading to myofiber breakdown^2,9^. Gain and loss of function FoxO mouse models have revealed that the FoxO transcription factors are key mediators of both the ubiquitin-proteasome and autophagy-mediated muscle proteolysis under diverse pathological conditions^10–13^. FoxO activates a transcriptional program that coordinates the loss of muscle mass, most often referred to as “atrogenes,” which include enzymes that catalyze autophagy, ubiquitin-proteasome activity, the unfolded protein response, mitochondrial function, and energy balance^14,15^. The most well studied “atrogenes” are two muscle specific ubiquitin ligases Atrogin-1/FBXO32 and MuRF1/TRIM63 that increase under catabolic conditions and ubiquitinate thick filament proteins leading to their proteasomal degradation^16–20^. Mice lacking Atrogin-1 or MuRF1 have partial resistance to muscle atrophy under multiple conditions, such as denervation, glucocorticoids, aging, and hindlimb unloading^21–24^. Atrogin-1 and MuRF1 are not required in the setting of starvation-or microgravity-mediated muscle atrophy nor are they sufficient to drive muscle atrophy suggesting that Atrogin-1- and MuRF1-independent pathways exist^10,16,18,20,25,26^. Conversely, FoxO regulates autophagy via control of several autophagy-related genes (*Atgs*)^2,9,12,27^. FoxOs are required to sustain autophagy under low nutrients, but FoxO inhibition does not affect basal autophagic flux^2^.

How low oxygen promotes muscle loss, such as in the context of high altitude, COPD, and PAH, remains unknown but involves accelerated proteolysis, metabolic derangement, and autophagy^4,28–30^. The HIF transcription factor, which consists of a labile α subunit (HIFα) and a stable β subunit (HIFβ/ARNT [aryl hydrocarbon nuclear translocator 1]), accumulates during hypoxia and activates genes that coordinate the cellular adaptation to low oxygen^3^. When oxygen is present HIFa becomes hydroxylated on one (or both) conserved proline residues by members of the Prolyl Hydroxylase Domain (PHD1-3, also known as the Egln-Nine [EglN]) family. PHD enzymatic activity requires oxygen, reduced iron, and 2-oxoglutarate and is sensitive to other upstream inputs, such as reactive oxygen species (ROS), and Krebs cycle intermediates^31,32^. Thus, the PHD proteins act as oxygen sensors to control HIFa. Once HIFa is hydroxylated, HIFa is polyubiquitinated by an E3 ubiquitin ligase complex containing pVHL (von Hippel-Lindau protein) leading to its proteasomal degradation^33^.

In the skeletal muscle, PHD2- or PHD1-deficient mice show loss of muscle mass and decreased muscle fiber size^34^. PHD1 loss in mice also reduces exercise capacity and decreased mitochondrial oxygen consumption^35^. Conversely, skeletal muscle PHD3 loss increases fatty acid oxidation and exercise capacity independent of HIF^36^. Skeletal muscle ARNT loss, which inactivates HIF, reduces skeletal muscle regeneration in aged mice^37^. Collectively, hypoxia and HIF signaling play a complex role in skeletal muscle homeostasis. Whether HIF plays an active, passive, or homeostatic role in hypoxia-induced muscle loss remains unclear. Further, whether HIF activates a similar “atrogene” program as FoxO in the setting of hypoxia-induced muscle atrophy remains unknown.

Here, we validate that either fasting or hypoxia is sufficient to induce muscle atrophy in mice. HIF is necessary and sufficient to mediate muscle atrophy under hypoxia. Transcriptional profiling of skeletal muscle isolated mice that are starved or housed under hypoxia identifies Decidual Protein Induced by Progesterone 1 (Depp1), which is activated under these conditions via FoxO1 and HIF, respectively. Using a combination of whole body Depp1 knockout mice and skeletal muscle-specific Depp1 transgenic animals, we find Depp1 is necessary and sufficient to induce skeletal muscle atrophy in the setting of fasting and hypoxia. Mechanistically, we find Depp1 is localized to the mitochondria and is necessary and sufficient to activate autophagy and mitochondrial degradation. Collectively, these studies nominate Depp1 as a critical “atrogene.”

## Results

### Fasting and hypoxia are sufficient to reduce muscle mass in mice

Nutrient deprivation is an established model of muscle atrophy, and the FoxO family of transcription factors are necessary and sufficient to mediate muscle loss under this condition^2^. On the other hand, how hypoxia leads to muscle loss is far less understood. To address this, we first asked whether we could detect hypoxia-induced muscle loss in vivo. We housed mice at 10% O_2_ (hypoxia) for 7-days and then euthanized these animals and measured muscle mass. As a positive control that we could detect muscle loss, we deprived an independent cohort of mice of food (fasted) for 48 hours. Food deprivation or hypoxia reduced gastrocnemius (GA) muscle mass and muscle fiber cross-sectional area (CSA) when compared to ad libitum fed mice housed at atmospheric oxygen (Figure 1A-B).

**Figure 1:**
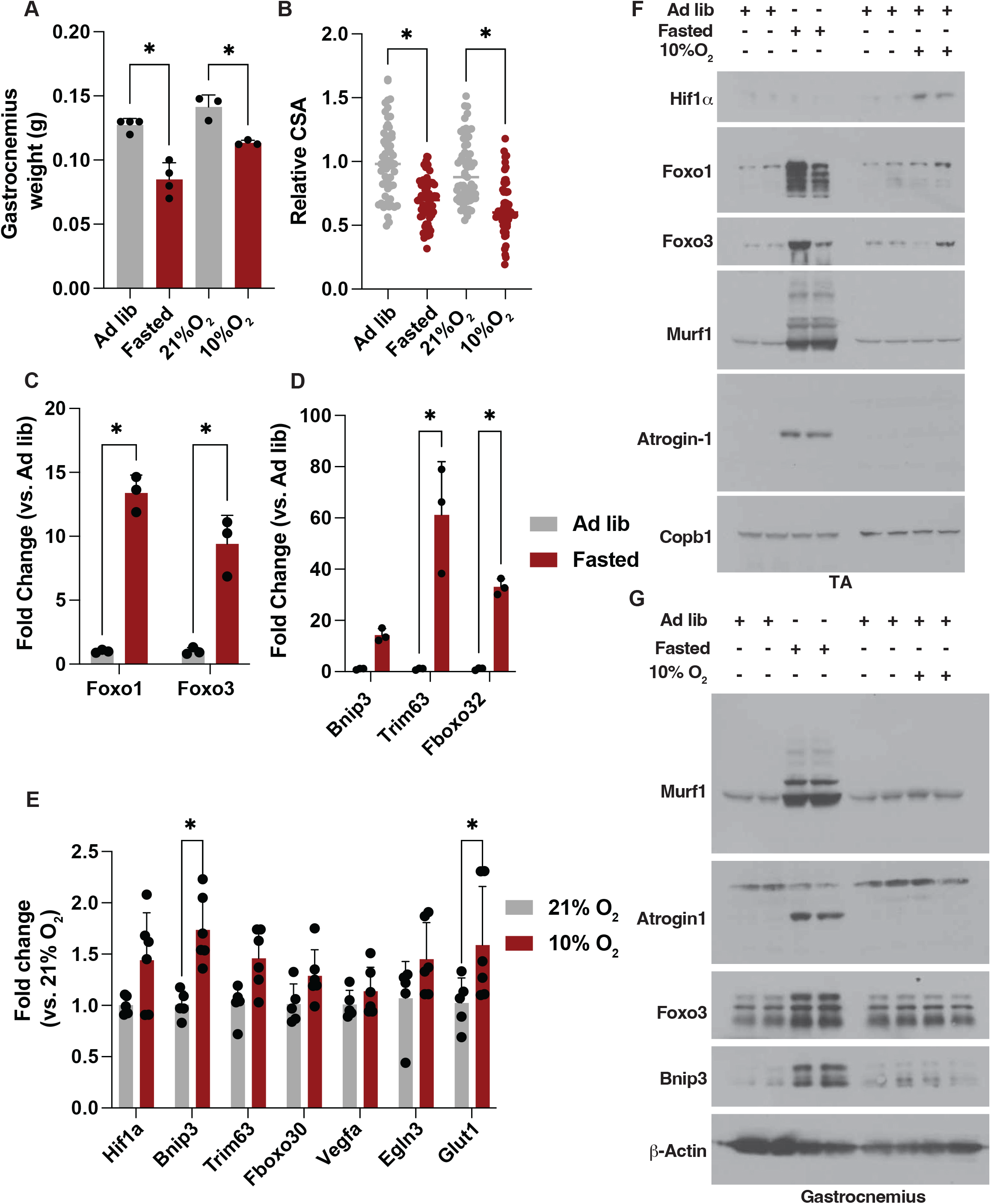
Hypoxia or fasting reduces muscle mass in mice. **(A-B)** Gastrocnemius muscle weight (grams, g) (A) and relative myofiber cross sectional area (CSA) (B) in mice. Where indicated, mice were fasted for 48 hours, housed at 10% O_2_ for 7 days, or fed ad libitum and housed at normal atmospheric oxygen (21% O_2_). * indicates P<0.05. N>3 per group. **(C-E)** mRNA analysis in gastrocnemius muscles isolated from mice deprived of food for 48 hours (fasted) (C-D), housed at 10% O_2_ for 7 days (E), or fed ad libitum and housed at normal atmospheric oxygen (21% O_2_). * indicates P<0.05. N=3 per group. **(F-G)** Immunoblot analysis of tibialis anterior (TA) (F) or gastrocnemius (G) muscle lysates isolated from mice deprived of food for 48 hours (fasted), housed at 10% O_2_ for 7 days, or fed ad libitum and housed at normal atmospheric oxygen (21% O_2_). Each sample is an individual mouse.

Nutrients and oxygen are sensed via the FoxO and HIF transcription factors, respectively, to control downstream gene expression^33,38^. We validated fasting or hypoxia activated FoxO and HIF, respectively, in GA and tibialis anterior (TA) muscles using quantitative polymerase chain reaction (qPCR) and immunoblot analysis (Figure 1C-G). Fasting increased Foxo1/3 RNA and protein abundance and increased downstream FoxO target gene expression, such as Bcl-2 interacting protein 3 (Bnip3), Trim63/MuRF1, Fboxo32/Atrogin-1, Unc-51 like autophagy activating kinase 1 (Ulk1), Autophagy related 14 (Atg14), and Cathepsin L (Figure 1C-D, F-G, and S1A-C). Hypoxia increased HIF abundance in TA muscle lysates via immunoblot analysis, and increased HIF-target gene expression, such as Bnip3 and Glut1, in GA muscles (Figure 1E-F). Despite an increase in Bnip3 RNA in hypoxic GA muscles, we did not detect an increase in Bnip3 protein abundance in this condition in GA muscle tissue lysates (Figure 1E and 1G). Fasting and hypoxia increased autophagy activation to a similar degree in GA muscle lysates, as measured by increased LC3B lipidation, but hypoxia did not promote the expression of autophagy-related genes that are known FoxO targets, such as Ulk1, Atg14, or Cathepsin L (Figure S1A-C and S2A-B). In hypoxic muscles, we did not detect an increase in FoxO1/3 abundance via qPCR or immunoblot analysis, nor did we detect an increase in Trim63/MuRF1 or Fboxo32/Atrogin-1 abundance in these assays despite a similar loss of muscle mass (Figure 1E-G). These results suggest hypoxia is sufficient to promote muscle loss and does so in a manner that is independent of FoxO, Trim63/MuRF1, or Fboxo32/Atrogin-1 activation.

### HIF activation is necessary and sufficient for hypoxia-induced muscle atrophy

HIF is the master regulator of genes that mediate the cellular adaptation to low oxygen^3^. Whether HIF is either necessary or sufficient to induce muscle atrophy under hypoxia remains unknown. To test whether HIF is necessary for hypoxia-induced muscle atrophy, we inactivated Arnt (HIFb), which is required for HIFa-mediated transcription, in mouse skeletal muscle using Arnt^fl/fl^ mice^39^. We crossed Arnt^fl/fl^ mice to skeletal muscle specific transgenic mice that express Cre recombinase driven by the human a-skeletal actin (*Acta*-Cre) promoter to generate Arnt^+/+^ Acta-Cre (wild-type) or Arnt^fl/fl^ Acta-Cre mice^40^. We confirmed Cre-mediated loss of Arnt in mouse GA muscle via immunoblot analysis (Figure 2A). We housed Arnt^+/+^ Acta-Cre and Arnt^fl/fl^ Acta-Cre under 10% O_2_ or normal atmospheric oxygen for 7-days prior to euthanasia. Hypoxia reduced GA muscle mass and GA CSA in wild-type mice, and this was lost in Arnt^fl/fl^ Acta-Cre mice (Figure 2B-C, S3A). Hypoxia did not induce Murf1 abundance in Arnt^+/+^ Acta-Cre and Arnt^fl/fl^ Acta-Cre GA muscle lysates (Figure 2A). Thus, HIF is necessary for hypoxia-induced muscle atrophy.

**Figure 2:**
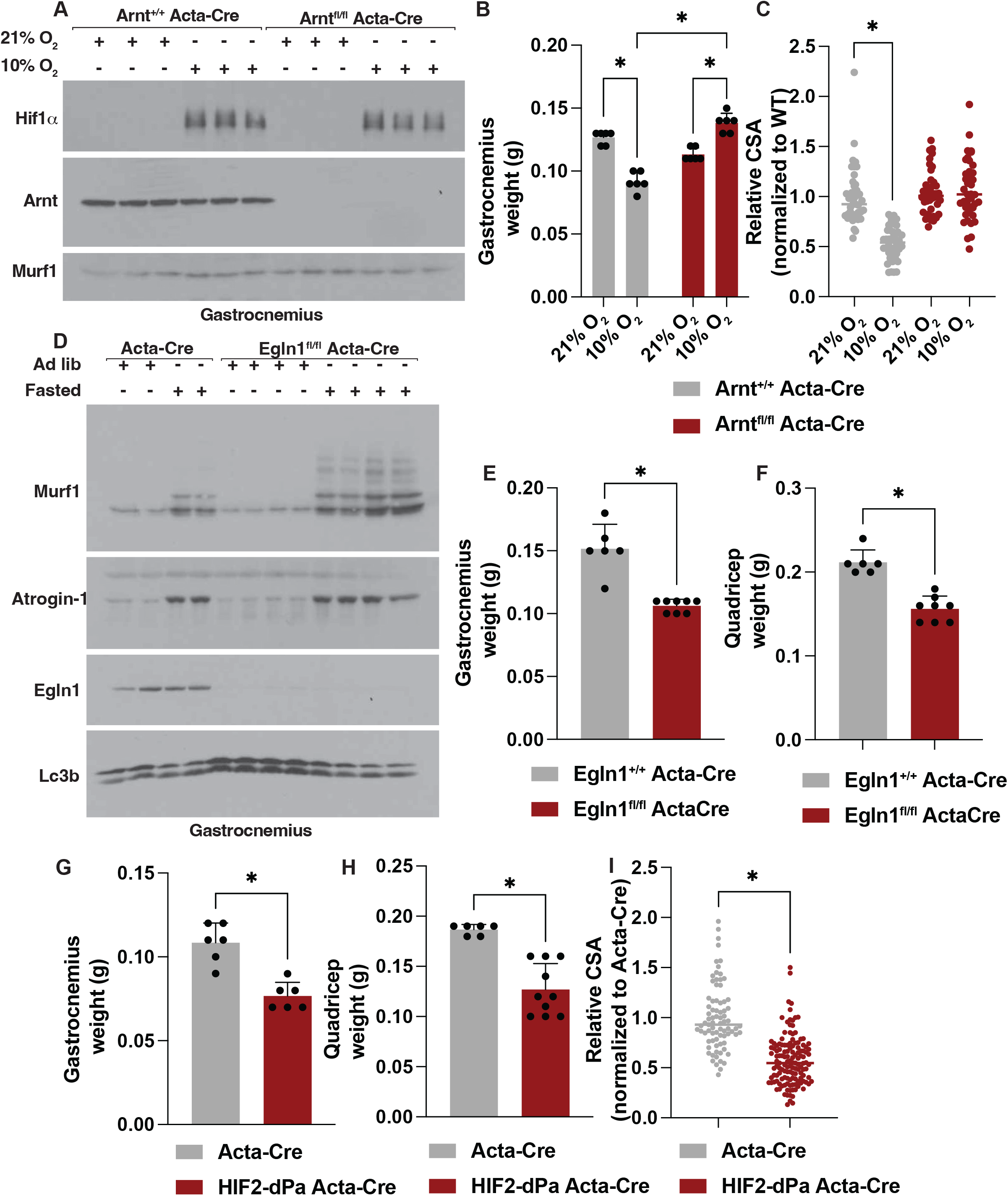
HIF is necessary and sufficient to mediate muscle loss under hypoxia in mice. **(A)** Immunoblot analysis of gastrocnemius muscle lysates isolated from Arnt^+/+^ Acta-Cre (Wild-type) and Arnt^fl/fl^ Acta-Cre mice. Where indicated, mice were housed at 10% O_2_ or normal atmospheric oxygen (21% O_2_) for 7 days prior to euthanasia. Each sample is an individual mouse. **(B-C)** Gastrocnemius muscle weight (grams, g) (B) and relative myofiber cross sectional area (CSA) (C) from Arnt^+/+^ Acta-Cre (Wild-type) and Arnt^fl/fl^ Acta-Cre mice. Where indicated, mice were housed at 10% O_2_ or normal atmospheric oxygen (21% O_2_) for 7 days prior to euthanasia. *indicates P<0.05. N>3 per group. **(D)** Immunoblot analysis of gastrocnemius muscle lysates isolated from Egln1^+/+^ Acta-Cre (Wild-type) and Elgn1^fl/fl^ Acta-Cre mice. Where indicated, mice were fasted for 48 hours or fed ad libitum prior to euthanasia. Each sample is an individual mouse. **(E-F)** Gastrocnemius (E) and quadricep (F) muscle weight (grams, g) from Egln1^+/+^ Acta-Cre (Wild-type) and Egln1^fl/fl^ Acta-Cre mice. *indicates P<0.05. N>3 per group. **(G-I)** Gastrocnemius (G) and quadricep (H) muscle weight (grams, g) and gastrocnemius myofiber cross sectional area (CSA) (I) from Acta-Cre (Wild-type) and HIF2-dPa Acta-Cre mice. *indicates P<0.05. N>3 per group.

To determine if HIF activation is sufficient to induce muscle atrophy, we first inactivated Egln1/PHD2, the primary HIFa hydroxylase in vivo, in mouse skeletal muscle^41–43^. We crossed Egln1^fl/fl^ mice to the skeletal muscle-specific Acta-Cre mice to generate Egln1^+/+^ Acta-Cre (wild-type) or Egln1^fl/fl^ Acta-Cre mice. We confirmed Cre-mediated loss of Egln1/PHD2 in mouse GA muscle via immunoblot analysis and measured GA and quadricep weights in these animals (Figure 2D). Egln1 loss reduced GA and quadricep mass when compared to wild-type animals (Figure 2E-F). Despite a decrease in muscle mass, Egln1 loss did not increase Trim63/MuRF1 or Fboxo32/Atrogin-1 RNA or protein abundance in GA muscle at baseline via qPCR and immunoblot analysis, suggesting muscle atrophy in the setting of Egln1 loss is not mediated via Trim63/MuRF1 or Fboxo32/Atrogin-1 activation similar to our results in hypoxic mice (Figure 2D and S3B). We confirmed we could detect an increase in Trim63/MuRF1 or Fboxo32/Atrogin-1 abundance in these mice upon FoxO activation via fasting (Figure 2D). Fasting increased Trim63/MuRF1 or Fboxo32/Atrogin-1 abundance in mouse GA muscle lysates isolated from wild-type and Egln1^+/+^ Acta-Cre (wild-type) and Egln1^fl/fl^ Acta-Cre mice (Figure 2D). Egln1 loss led to an increased abundance of Trim63/MuRF1 in GA muscle lysates upon fasting, possibly due to these muscles being “primed” for atrophy via a different stimulus (Figure 2D).

As Egln1 has HIF-independent functions, we tested whether HIF activation alone is sufficient to promote muscle loss. To test this, we crossed Acta-Cre mice with mice that carry a conditional HIF2α allele that contains a LoxP-Stop-LoxP cassette upstream of a cDNA encoding a non-hydroxylatable HIF2α (HIF2α dPA), which escapes recognition by the VHL E3 ligase and is sufficient to activate HIF signaling in the presence of oxygen^44^. As HIF2α dPA in the heart leads to myocyte loss, we predicted it would be sufficient to reduce skeletal muscle mass when expressed in a skeletal muscle specific manner^45^. Indeed, HIF2α dPA reduced GA and quadricep muscle mass and GA myofiber cross sectional area (CSA) when compared to Acta-Cre mice (Figure 2G-I and S3C). Collectively, these results support HIF is necessary and sufficient to promote hypoxia-induced muscle atrophy, and this is independent of Trim63/MuRF1 or Fboxo32/Atrogin-1 activation.

### Transcriptional profiling identifies Depp1 in starved or hypoxic muscle

The muscle specific E3 ubiquitin ligases Trim63/MuRF1 and Fboxo32/Atrogin-1 are the most validated mediators of FoxO-induced muscle atrophy, however they are not induced under hypoxia suggesting other unknown “atrogenes” remain to be discovered.

As HIF is sufficient to drive muscle atrophy in vivo, we hypothesized HIF may control an unidentified target gene(s) that mediates skeletal muscle atrophy under hypoxia^13,27^. As FoxO is also sufficient to drive muscle atrophy in vivo, it is possible FoxO and HIF may control an unidentified overlapping target gene(s) that mediates skeletal muscle atrophy. To address this, we utilized RNA sequencing in mouse GA muscles isolated from wild-type Acta-Cre animals that were housed under 10% O_2_ for 24 hours to activate HIF or fasted for 48 hours to activate FoxO (Figure 3A-C). To determine genes that are controlled via the HIF transcription factor under hypoxia in the skeletal muscle, we also housed Arnt^fl/fl^ Acta-Cre under hypoxia. As expected, hypoxia increased the expression of known HIF-target genes in the GA muscle, such *Bnip3*, *Vegfd*, *Ndrg1,* which were suppressed in GA muscles lacking Arnt (Figure 3A-B). Hypoxia elevated *Foxo1*, *Foxo3*, *Trim63*/MuRF1, and *Fboxo32*/Atrogin-1 RNA abundance and this was unaffected by Arnt loss, although we did not detect an increase in the protein products of these genes in our immunoblot experiments (Figure 3A-B and 1F-G). As expected, fasting increased the expression of *Foxo1* and *Foxo3* and downstream FoxO target genes, such as *Trim63*/MuRF1, *Fboxo32*/Atrogin-1, *Gabarapl1,* and *Map1lc3b* (Figure 3C). Among the hypoxia and fasting-induced genes, we identified Decidual Protein Induced by Progesterone 1 (Depp1) (Figure 3A-C). Depp1 was of interest as its role in the context of skeletal muscle atrophy has not been studied but it has been shown to be upregulated in multiple mouse models of muscle atrophy, such as aging, microgravity, and fasting^27,46,47^. Arnt loss blocked the hypoxia-regulated elevation of *Depp1*, supporting the hypothesis that HIF controls Depp1 in skeletal muscle (Figure 3A-B). Similarly, Depp1 is elevated in *Trim63*/MuRF1 knockout mouse muscles, which are partially resistant to atrophy across multiple models of FoxO activation^23^. In the context of fasting, Depp1 expression is controlled via FoxO1 in C2C12 mouse myoblast cells^27^. We have previously shown that it plays a key role in HIF-mediated cardiac remodeling in the setting of heart failure^48^. Similarly, DEPP1 has been linked with autophagy activation via FoxO before in neuroblastoma cells^49^. We validated Depp1 is increased in mouse GA muscles upon fasting and hypoxia via qPCR (Figure S4A-B). Similarly, we validated Depp1 is increased in the skeletal muscle of another FoxO-mediated atrophy model, specifically in the setting of dexamethasone treatment (Figure S4C)^20,23^. We have been unable to identify a commercially available Depp1 antibody that detects Depp1 in mouse tissues. To circumvent this problem, we utilized Crispr-Cas9 gene editing in mouse embryos to modify the Depp1 locus such that it encodes for an in-frame C-terminal hemagglutinin (HA) tag to allow us to detect Depp1 protein in mouse skeletal muscle in immunoblot experiments using an anti-HA antibody. Like our qPCR results, fasting and hypoxia increased Depp1 protein abundance in vivo (Figure S4D).

**Figure 3:**
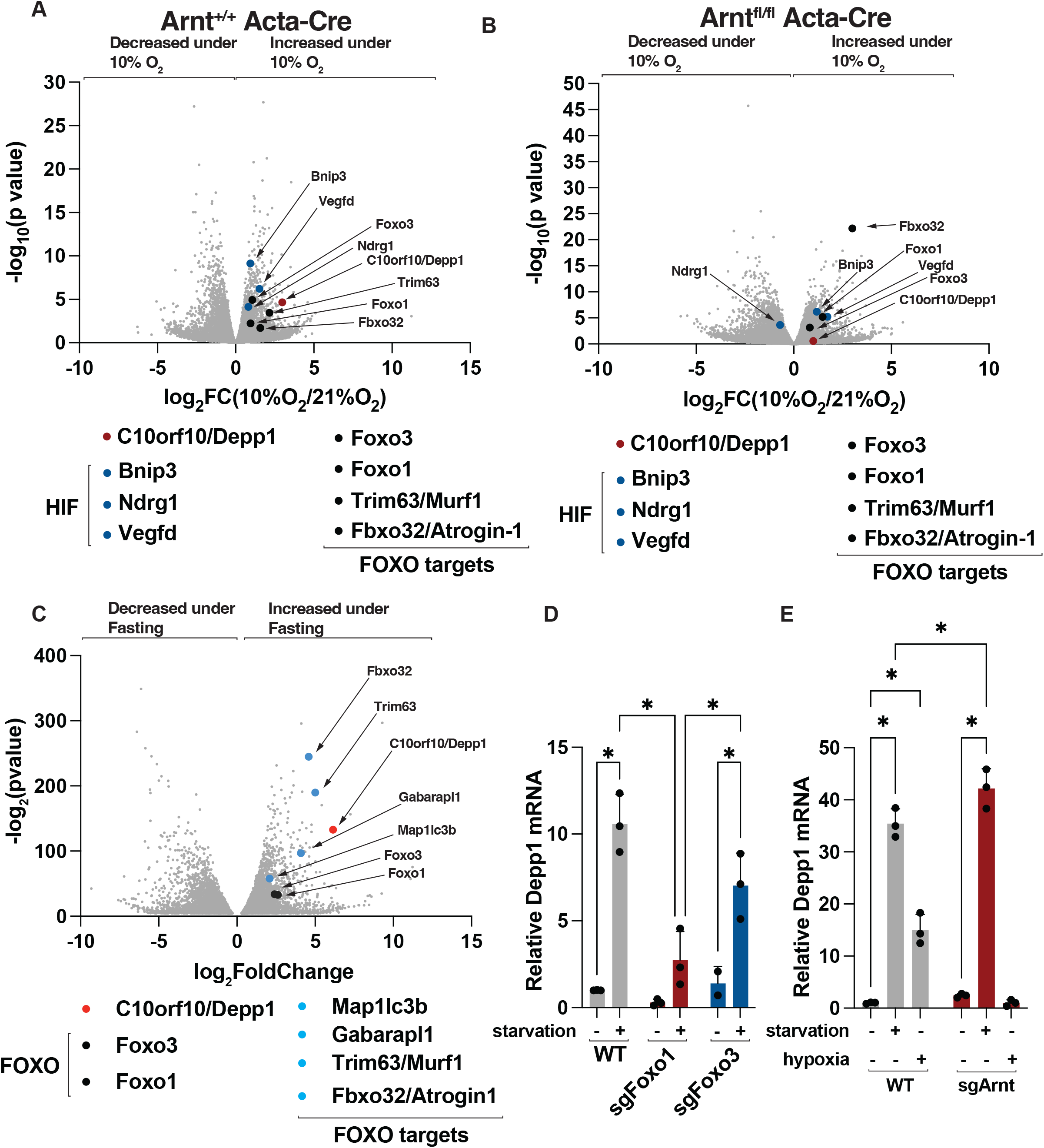
Depp1 is elevated in the skeletal muscle under hypoxia or fasting. **(A-C)** Transcriptomic profiling of RNA isolated from Acta-Cre (Wild-type) and Arnt^fl/fl^ Acta-Cre mice gastrocnemius muscles. Where indicated, mice were housed at 10% O_2_ for 7 days (A-B), deprived of food for 48 hours (fasted) (C), or fed ad libitum at normal atmospheric oxygen (21% O_2_) prior to euthanasia. N=3 per group. (D-E) mRNA analysis in wild-type, sgFoxo1, sgFoxo3 (D), and sgArnt (E) C2C12 cells. Where indicated, cells were deprived of nutrients (starvation) (D-E), grown at 0.1% O_2_ (hypoxia), or grown in full media at normal atmospheric oxygen for 18 hours prior to RNA extraction. * indicates P<0.05. N=3 per group.

To confirm the relevant transcription factors that control Depp1 expression upon fasting and hypoxia, we generated C2C12 cells lacking either Foxo1, Foxo3, or Arnt using CRISPR-Cas9 (Figure S5A-C). Nutrient starvation increased Foxo1 and Foxo3 abundance in immunoblot assays (Figure S5A). We detected a compensatory increase in Foxo1 or Foxo3 abundance in nutrient starved cells where the other FoxO family member is inhibited (Figure S5A). Similarly, Foxo1 or Foxo3 loss increased HIF stabilization under hypoxia suggesting FoxO suppresses HIF activation, consistent with prior studies (Figure S5A)^50^. We confirmed Arnt loss blocked HIF-transcription as measured by reduced stabilization of HIF-target gene Bnip3, under hypoxia (Figure S5B-C). Nutrient starvation increased *Depp1* RNA and this required Foxo1, but not Foxo3 (Figure 3D). Conversely, Arnt loss in C2C12 cells blocked the hypoxia-mediated increased in *Depp1* RNA, consistent with our RNA sequencing results *in vivo*, but did not suppress the increase in *Depp1* upon nutrient starvation (Figure 3E). Collectively, Depp1 is induced in mouse muscles under conditions of fasting and hypoxia and this is mediated via distinct transcriptional pathways involving Foxo1 and HIF, respectively.

### Depp1 is necessary and sufficient to mediate muscle loss under fasting and hypoxia

To test the role of Depp1 in fasting- and hypoxia-induced muscle atrophy, we utilized whole body *Depp1^-/-^* mice^51^. Our previous studies have shown *Depp1^-/-^* mice are viable, normotensive, and do display gross phenotypes, consistent with previous reports of similar mice^48,52,53^. Food deprivation for 48 hours or 7-days hypoxia (10%O_2_) reduced GA mass and cross-sectional area (CSA) in wild-type mice, and this effect was blunted in *Depp1^-/-^* mice (Figure 4A-F). Muscle atrophy results in a loss of myofibrillar proteins, such as fast and slow twitch myosin heavy chain (MyHC)^18,20^. As Depp1 loss protected against muscle atrophy, we analyzed muscle lysates from fasted Depp1^-/-^ mice to identify proteins that were preserved. We confirmed fasting activated FoxO signaling in wild-type and *Depp1*^-/-^ mouse TA and GA muscles as measured by increased MurF1, Atrogin-1, Bnip3, and FoxO1 abundance in immunoblot and qPCR experiments (Figure S6A-D). The elevation in FoxO target gene expression upon fasting was increased in *Depp1^-/-^* mice over fasted wild-type mice in qPCR experiments, however, where tested, this was not reflected at the protein level in immunoblot experiments (Figure S6D). Fasting reduced fast and slow twitch MyHC abundance in wild-type mouse TA muscle lysates when compared to *ad libitum* fed mice (Figure S6D-F). In absence of Depp1, MyHC protein abundance was preserved in TA muscle lysates isolated from fasted mice when compared to *ad libitum* fed animals (Figure S6D-F).

**Figure 4:**
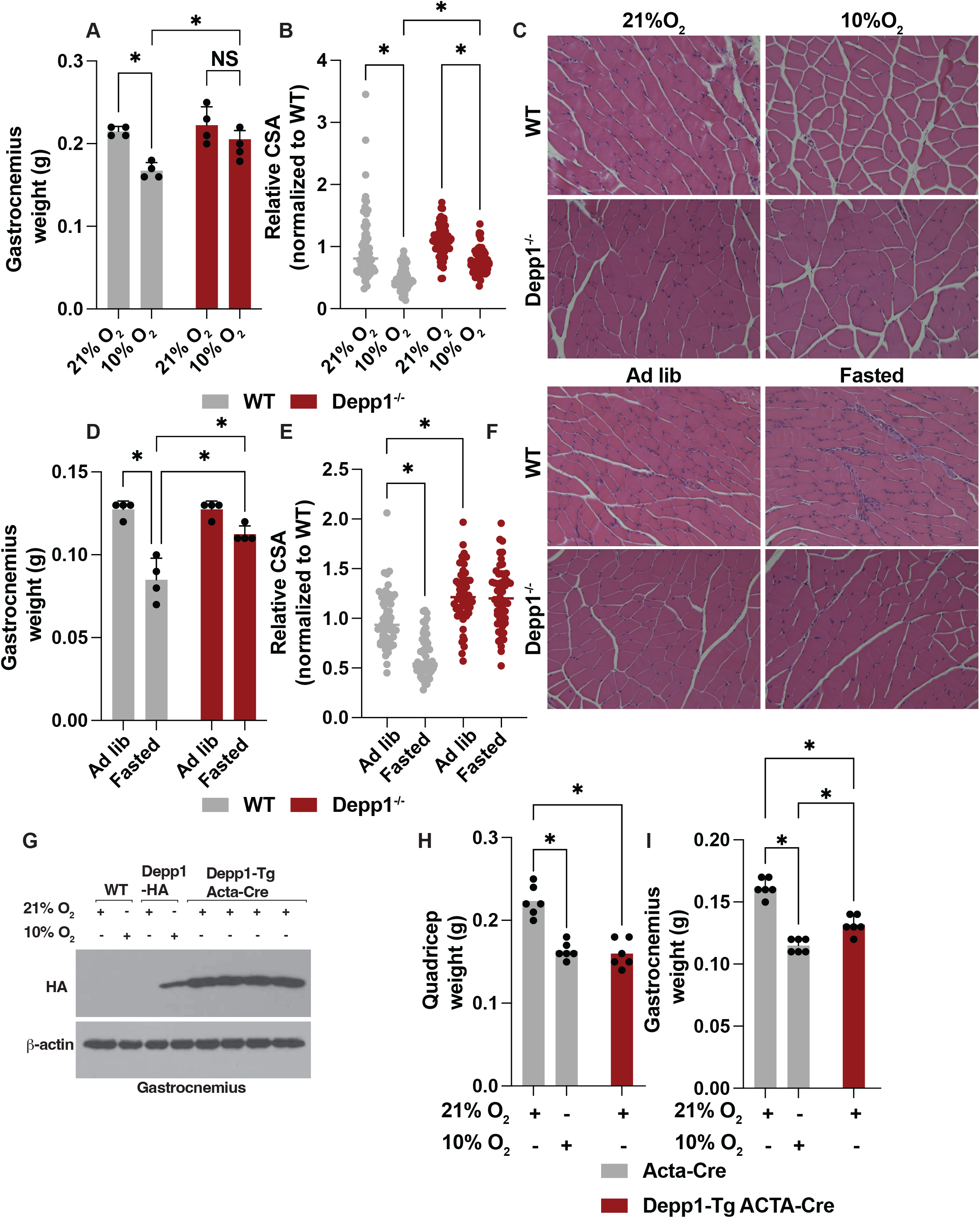
Depp1 is necessary and sufficient to mediate muscle atrophy under fasting and hypoxia. **(A-C)** Gastrocnemius muscle weight (grams, g) (A) and relative myofiber cross sectional area (CSA) (B) and gastrocnemius Hematoxylin and Eosin (HE) staining (C) from wild-type (WT) and Depp1^-/-^ mice. Where indicated, mice were housed at 10% O_2_ or normal atmospheric oxygen (21% O_2_) for 7 days prior to euthanasia. *indicates P<0.05. N>3 per group. **(D-F)** Gastrocnemius muscle weight (grams, g) (D) and relative myofiber cross sectional area (CSA) (E) and gastrocnemius Hematoxylin and Eosin (HE) staining (R) from wild-type (WT) and Depp1^-/-^ mice. Where indicated, mice were deprived of food (fasting) for 48 hours or fed ad libitum prior to euthanasia. *indicates P<0.05. N>3 per group. **(G)** Immunoblot analysis of gastrocnemius muscle lysates isolated from wild-type (WT), endogenous Depp1-HA knock-in, and Depp1-Tg Acta-Cre mice. Where indicated, mice were housed at 10% O_2_ or normal atmospheric oxygen (21% O_2_) for 7 days prior to euthanasia. **(H-I)** Quadricep (H) and gastrocnemius (I) muscle weight (grams, g) from Acta-Cre (WT) and Depp1-Tg Acta-Cre mice. Where indicated, mice were housed at 10% O_2_ or normal atmospheric oxygen (21% O_2_) for 7 days prior to euthanasia. *indicates P<0.05. N>3 per group.

Increased Vascular Endothelial Growth Factor A (Vegfa) preserves muscle mass in aging and injury models via increased vascularization and muscle fiber regeneration^54,55^. Vegfa is also regulated via HIF in many cell types^3,56^. While we did not detect a hypoxia-mediated increase of *Vegfa* mRNA in gastrocnemius muscles isolated from wild-type mice, Depp1 loss induced *Vegfa* mRNA abundance under hypoxia suggesting Depp1 inhibition may stimulate Vegfa to preserve muscle mass under this condition (Figure S7A).

To test whether Depp1 is sufficient to promote muscle loss, we generated a conditional Depp1 transgenic mouse (Depp1-Tg) in which the mouse Depp1 complementary DNA (cDNA) sequence fused to a c-terminal HA tag is knocked into the ROSA26 safe harbor locus downstream of a LoxP flanked neomycin (Neo) resistance stop selection marker and a cytomegalovirus (CMV) early enhancer/chicken b actin (pCAG) promoter. We crossed our Depp1-Tg mice to skeletal muscle specific Acta-Cre mice to generate Depp1-Tg Acta-Cre mice. We validated increased Depp1 expression in Depp1-Tg Acta-Cre TA tissue lysates isolated from 3-week-old mice when compared against Acta-Cre TA muscles via anti-HA immunoblotting experiments (Figure 4G). Exogenous Depp1 expression in our Depp1-Tg Acta-Cre mice was roughly four-fold higher than endogenous Depp1 abundance in mouse TA lysates isolated from our endogenously tagged Depp1-HA mice housed at 10% O_2_ for 7-days (Figure 4G). We euthanized the Depp1-Tg animals at 4 months of age and measured muscle mass. Depp1-Tg reduced GA and quadricep muscle weight when compared to Acta-Cre mice (Figure 4H-I). Collectively, these data support that Depp1 is both necessary and sufficient to induce muscle atrophy *in vivo*.

### Depp1 is necessary to activate autophagy under fasting and hypoxia in vivo

Autophagy activation is a common downstream consequence of FoxO and HIF activation and is important to drive muscle atrophy^9,57^. The precise mechanism by which the FoxO family activates autophagy remains unclear, but FoxO is required to sustain autophagic flux under low nutrients in the muscle but does not affect basal autophagic flux^2,9^. We have previously shown that Depp1 is necessary and sufficient to control HIF-mediated autophagy in the heart^48^. To test whether Depp1 is necessary for fasting or hypoxia-induced autophagy in skeletal muscle, we utilized the transgenic *CAG-RFP-GFP-LC3* mouse line in which a CAG promoter sequence drives the expression of red fluorescent protein (RFP) and an enhanced green fluorescent protein (EGFP) fused to microtubule-associated protein 1 light chain 3 alpha (LC3)^58,59^. Upon autophagy activation, the *RFP-GFP-LC3B* fusion protein is trafficked to the lysosome and visualized via confocal microscopy where the EGFP signal is quenched while the RFP protein accumulates^59^. An increase in RFP puncta is indicative of increased autophagy activation. We crossed these mice to our *Depp1^-/-^* mice and measured autophagy activation in mouse gastrocnemius muscles following fasting or hypoxia (Figure 5A-D). Fasting or hypoxia increased RFP puncta in wild-type mouse gastrocnemius muscles and this was reduced in mouse gastrocnemius muscles isolated from *Depp1^-/-^* mice (Figure 5A-D). In line with these results, where tested, Depp1 loss reduced Lc3b lipidation in mouse quadriceps in immunoblot experiments (Figure S8A-B).

**Figure 5:**
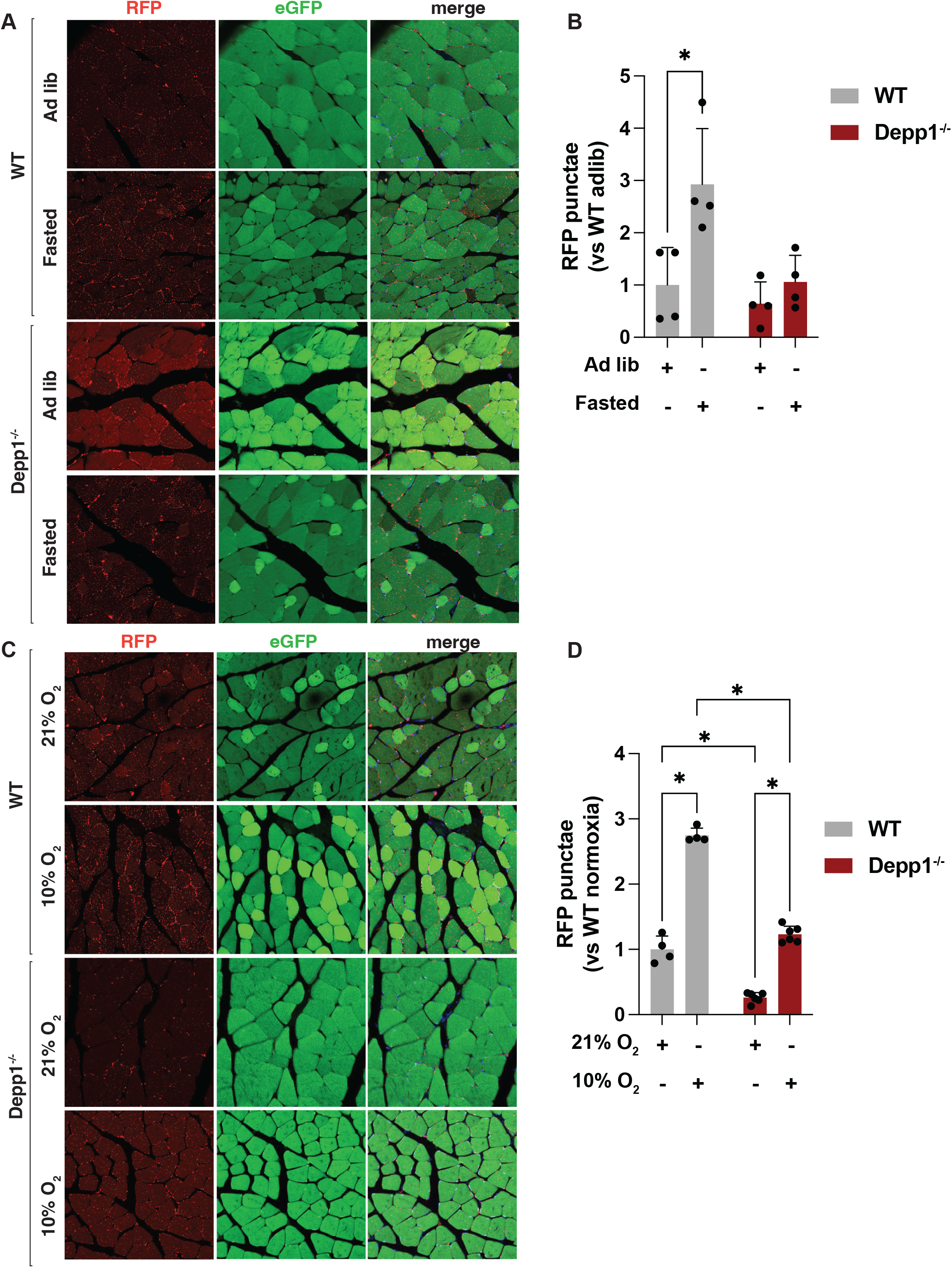
Depp1 is necessary to activate autophagy in the skeletal muscle under hypoxia and fasting. **(A-D)** Confocal microscopy and image analysis of RFP-eGFP-Lc3 wild-type (WT) and Depp1^-/-^ mouse gastrocnemius muscles. Where indicated, mice were deprived of food (fasted) for 48 hours or fed ad libitum (A-B), housed at 10% O_2_ or fed ad libitum and housed at normal atmospheric oxygen (21% O_2_) (C-D) for 7 days prior to euthanasia. *indicates P<0.05. N>3 per group.

### Depp1 loss preserves mitochondrial abundance under fasting and hypoxia

We have previously shown Depp1 localizes inside the mitochondria in cardiomyocytes, and this requires its N-terminal t-snare domain (Tsnare)^48^. To test whether this is true in skeletal muscle, we stably expressed Depp1 fused to mNeonGreen (Depp1-mNG) in human skeletal muscle cells. Like cardiomyocytes, Depp1 localized to mitochondria in human skeletal muscle cells as measured by its co-localization with MitoTracker Deep Red (Figure 6A). This was specific to the mitochondria as Depp1 did not localize to another organelle like the lysosome, as measured by LysoTracker Blue (Figure 6A). Lysosomes were found near Depp1-positive mitochondria consistent with lysosome-mitochondria contact sites^60^. As Depp1 is localized to the mitochondria, we hypothesized Depp1 regulates mitochondrial function. To test this, we measured mitochondrial oxygen consumption using Seahorse respirometry in C2C12 cells engineered to over-express Depp1. Using lentiviral transduction, we stably expressed DEPP1-mNG or empty vector (EV) in C2C12 cells. DEPP1 reduced mitochondrial oxygen consumption when compared to EV expressing C2C12 cells (Figure 6B).

**Figure 6:**
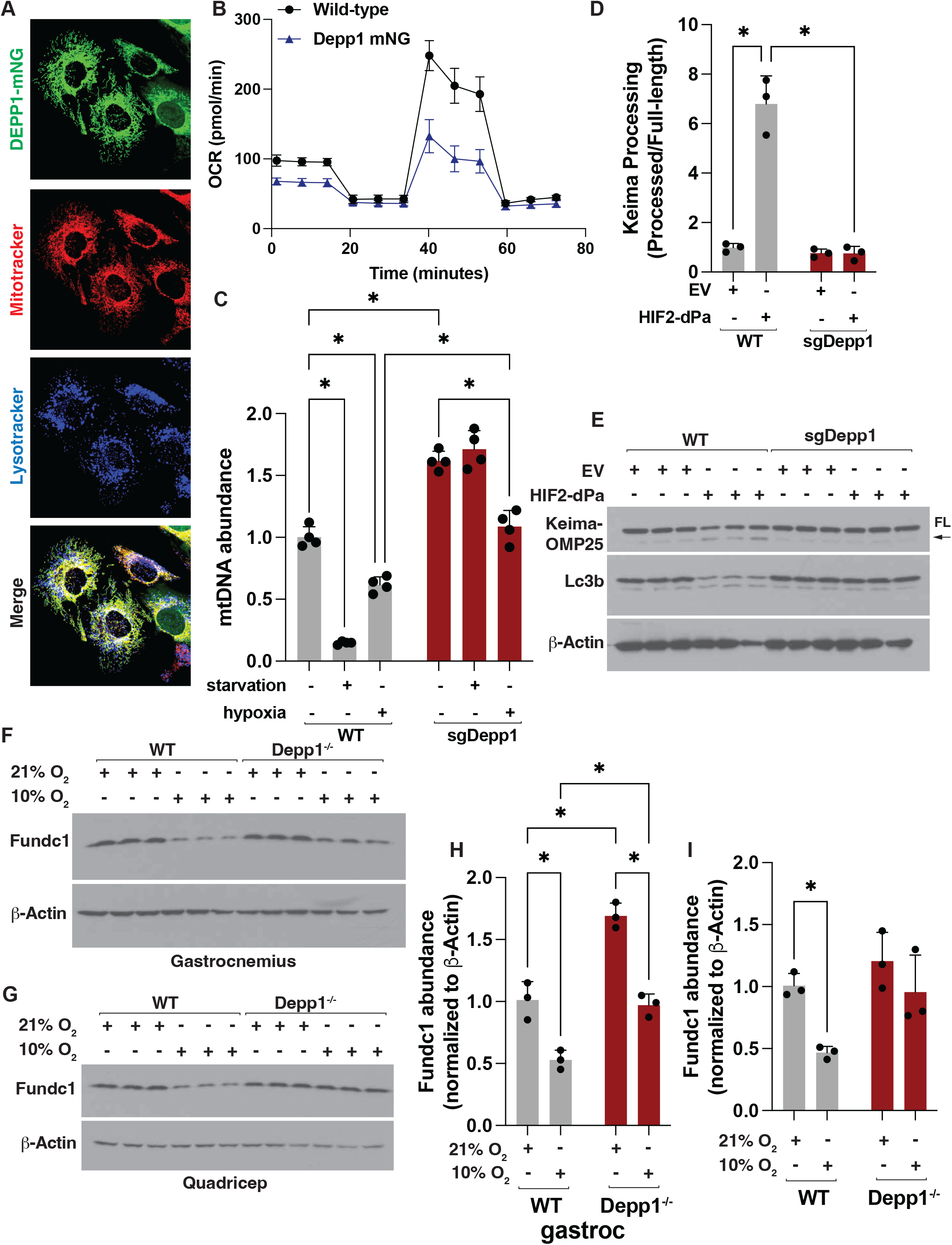
Depp1 is necessary for mitophagy in skeletal muscle. **(A)** Confocal microscopy analysis of human skeletal muscle cells stably expressing DEPP1 fused to mNeonGreen (DEPP1-mNG) and labeled with 500 nM MitoTracker DeepRed and 100 nM LysoTracker Blue. **(B)** Seahorse respirometry of C2C12 cells stably expressing DEPP1-mNG or empty vector (EV). **(C)** Mitochondrial DNA (mtDNA) abundance as measured by quantitative polymerase chain reaction (qPCR) in wild-type (WT) and sgDepp1 C2C12 cells. Where indicated, cells were deprived of nutrients (starvation), grown at 0.1% O_2_ (hypoxia), or grown at normal atmospheric oxygen in full media for 18 hours prior to RNA extraction. *indicates P<0.05. N>3 per group. **(D-E)** Densitometry (D) and immununoblot analysis (E) of wild-type (WT) and sgDepp1 C2C12 cells stably expressing Keima-OMP25. Where indicated, cells were also expressing HIF2-dPa or empty vector (EV). FL indicates full-length Keima-OMP25 fusion protein and arrow indicates processed Keima. **(F-I)** Immunoblot (F-G) and quantification (H-I) of mouse gastrocnemius (F, H) and quadricep (G, I) muscle lysates isolated from wild-type (WT) and Depp1^-/-^ mice. Where indicated, mice were housed at 10% O_2_ for 7 days or housed at normal atmospheric oxygen (21% O_2_). * indicates P<0.05. N=3 per group.

As Depp1 is necessary to activate autophagy under fasting and hypoxia in the skeletal muscle and localizes to the mitochondria, we hypothesized that Depp1 is necessary to degrade mitochondria (mitophagy) under conditions of HIF or FoxO activation. To test this, we generated *Depp1*^-/-^ C2C12 cells using CRISPR-Cas9 and measured mitochondria DNA (mtDNA), as a surrogate for mitochondria abundance, using qPCR under hypoxia and nutrient starvation (Figure S9A). Wild-type C2C12 cells grown under hypoxia or starved of nutrients increased Depp1 mRNA abundance and decreased mtDNA under these conditions (Figure S9A and 6C). Depp1 loss increased mtDNA abundance at baseline and preserved mtDNA abundance under conditions of nutrient starvation and hypoxia (Figure 6C). Bnip3 is an established hypoxia-mediated mitophagy receptor^61–64^. Like Depp1, hypoxia induced Bnip3 mRNA in wild-type C2C12 cells (Figure S9B). In cells lacking Depp1, Bnip3 mRNA abundance was elevated at baseline suggesting cells may compensate for loss of Depp1-mediated mitophagy via increasing Bnip3-mediated mitochondrial degradation (Figure S9B). To test whether Depp1 is important for mitophagy directly, we utilized our previously reported MITO-Keima reporter in which Keima is fused to the N-terminus of the classic, outer mitochondrial membrane localization sequence of OMP25 (Keima-OMP25)^48^. Keima fusion proteins facilitate the study of trafficking of autophagy cargo as the Keima protein is stable to lysosomal proteases, therefore when a Keima fusion protein enters the lysosome, it frees “processed” Keima (Keima and any peptide remnants left behind following proteolysis of its fusion partner in the lysosome), which can be detected via anti-Keima immunoblotting^65^. We stably expressed our Keima-OMP25 in wild-type and Depp1^-/-^ C2C12 cells using lentiviral transduction. Following, we stably expressed a cDNA encoding a non-hydroxylatable HIF2α (HIF2α dPA) that escapes recognition by the VHL E3 ligase, which we validated is sufficient to induce muscle atrophy in vivo or empty vector (EV)^44,66^. In wild-type C2C12 cells, HIF2α dPA increased Keima-OMP25 processing and this effect was lost in cells lacking Depp1 (Figure 6D-E). To determine whether Depp1 is important for mitophagy in vivo, we measured FUN14 domain-containing protein 1 (Fundc1) abundance in quadriceps and gastrocnemius muscles. Fundc1 is a key outer mitochondrial membrane protein that acts as a receptor for hypoxia-induced mitophagy; therefore, its abundance acts as a surrogate for mitophagy^67^. Hypoxia reduced Fundc1 abundance in mouse gastrocnemius muscles and this effect was HIF- and autophagy-dependent as the effect was lost in mouse gastrocnemius muscles lacking Arnt or Atg7 (Figure S10A-B). Atg7 loss in skeletal muscle increased Nbr1 abundance, a known autophagy receptor (S10A)^68,69^. In mice lacking Depp1, Fundc1 abundance was preserved under hypoxia in both quadriceps and gastrocnemius muscles when compared to wild-type (Figure 6F-I). Thus, Depp1 is a key factor in mitochondrial degradation in the muscle.

## Discussion

Nutrient availability, oxygen abundance, and mechanical and neuronal inputs, are essential to maintain muscle mass^70^. Disruption of any of these is sufficient to induce muscle atrophy, such as in the setting of chronic starvation, high altitude exposure, mechanical unloading, or neurodegeneration. Defining the mechanisms that lead to atrophy when any of these critical signals are lost, and whether these mechanisms are shared across diverse atrophy conditions, remains a central question to ultimately combat muscle loss to improve quality of life and health span. Here, we focus on the disruption of two of these critical upstream cues: oxygen and nutrients. We define a novel overlapping mechanism by which nutrient and oxygen abundance converge on Depp1 in the skeletal muscle via the FoxO and HIF transcription factors. Using gain- and loss-of-function mouse models, we find Depp1 is necessary and sufficient to promote muscle atrophy under conditions of starvation and hypoxia. Mechanistically, Depp1 localizes to the mitochondria where it is necessary to control autophagy activation and mitochondrial degradation. Thus, Depp1 is a previously uncharacterized “atrogene” critical for muscle atrophy across scenarios in which nutrients or oxygen is limiting.

While many studies have primarily focused on the FoxO family as critical mediators of muscle atrophy, we now report that HIF plays a critical role in the skeletal muscle to control an atrophy gene expression program in the muscle that is activated under low oxygen. We find HIF is necessary and sufficient to mediate muscle atrophy and this is independent of core “atrogene” factors Murf1/TRIM63 and Atrogin-1/FBXO32 activation. Instead, we find Depp1 is a critical mediator of muscle atrophy under low oxygen. Consistent with this, previous work has shown pharmacologic HIF2a inhibition using PT-2385 can ameliorate chronic hypoxia-associated muscle loss in mouse models via improving muscle regeneration^71^. While tissue hypoxia occurs in a multitude of pathological conditions, healthy individuals often experience low oxygen via transient or chronic exposure to high altitude. High altitude exposure is sufficient to drive body weight loss with muscle mass loss constituting upwards of 20% of the total weight lost. This is far more extreme in high-altitude athletes (above 14,000 feet) who often lose nearly 60% of muscle mass during high altitude exertion. Thus, HIF inhibition under these contexts may provide a strategy to mitigate muscle wasting. While our results on Depp1 focus on a cell-intrinsic mechanism of muscle atrophy in the context of HIF and FoxO activation, they do not preclude the existence of cell-extrinsic mechanisms that mediate muscle loss when nutrients or oxygen are limiting. Indeed, HIF2-driven clear cell carcinomas (ccRCC) promote Parathyroid hormone-related protein (PTHrP) secretion that is a critical driver of cancer-associated cachexia in this context^72^.

While our findings are consistent with earlier reports that fasting or low oxygen can induce Depp1 expression in skeletal muscle, starvation or hypoxia can affect skeletal muscle gene expression in many ways independent of HIF or FoxO, such as via Transcription factor EB (Tfeb), Activating Transcription Factor 4 (Atf4), or inhibiting histone demethylases^73–76^. Our studies conclude that nutrient or oxygen limitation induces Depp1 expression in the skeletal muscle via Foxo1 and HIF, respectively. This is consistent with prior work in the heart and gain- and loss-of-function FoxO family mouse models^27,48^. In addition, Depp1 mRNA is induced in tissues in aged mice and mice undergoing simulated microgravity^46,47^. Age-related muscle atrophy (sarcopenia) is strongly associated with frailty and increased risk of chronic disease^77,78^. While age-related muscle loss is mediated via multiple factors, such as decreased endocrine and neuronal activity, nutritional changes, or increased inactivity, the molecular mechanisms that contribute to age-related muscle loss remain to be elucidated^79^. Future studies will need to address whether Depp1 loss preserves muscle mass during aging. Similarly, microgravity-induced muscle loss is a common and debilitating consequence of long-duration space-flight missions and is a serious complicating factor in the possibility of humans becoming an interplanetary species^80,81^. How we lose muscle mass in low gravity conditions remains unknown but is independent of MuRF1/TRIM63 in mouse models^26^. Whether Depp1 loss preserves muscle mass following microgravity exposure or during spaceflight will need to be determined.

Skeletal muscle mitochondria make up roughly 6% of skeletal muscle fiber volume in humans^82,83^. As such, mitochondrial dysfunction is intimately linked to muscle wasting. Mitophagy is a quality control housekeeping program that maintains mitochondrial abundance and health under basal conditions and under stress via autophagy^84^. Chronic autophagy activation or inhibition both result in muscle atrophy and can exacerbate muscle wasting under stress^85,86^. Similarly, autophagy inhibition in muscle via Atg7 loss leads to the accumulation of abnormal mitochondria, decline in energy production, and muscle loss^85^. We find Depp1 is necessary for autophagy activation in the setting of hypoxia and starvation and preserves mitochondrial abundance under these conditions. Contrary to mouse models that completely lack autophagy, Depp1 loss does not negatively affect skeletal muscle homeostasis supporting that it does not impact basal autophagy flux, which is necessary to preserve mitochondrial function and maintain myofiber integrity. How Depp1 regulates mitochondria remains unclear, however mitophagy inhibition via inhibition of selective mitophagy receptor Bnip3, rather than the general autophagy machinery, has been shown to preserve mitochondrial homeostasis and reduces muscle loss in the setting of cancer cachexia mouse models or atrophy models mediated via Fibroblast growth factor 21 (FGF21)^87–89^. This appears context dependent as Bnip3 overexpression preserves muscle homeostasis during aging in *Drosophila*^90^.

The biochemical function of Depp1 remains to be determined. In our results in skeletal muscle and the heart, Depp1 plays a critical role in mitophagy in the context of hypoxia supporting that this is a conserved Depp1 function across tissues^48^. Here, we extend those findings and identify Depp1 is a key mitophagy factor when nutrients are limiting in the skeletal muscle. Depp1 is not a mitophagy receptor in the same vein as Fundc1 or Bnip3 as it is imported into the mitochondria and does not directly interface with the autophagy machinery in the cytoplasm, nor have we detected any interaction of Depp1 with ATG8-family proteins in biochemical assays^48^. We and others have shown Depp1 is sufficient to activate autophagy *in vivo* in multiple contexts^48,49,53,91,92^. As Depp1 localizes to the mitochondria and when lost reduces autophagy activation *in vivo* and preserves mitochondrial abundance, we predict Depp1 signals an “inside-out” mechanism to regulate autophagy activation and mitochondrial degradation that will need to be determined.

## Acknowledgements

We thank the members of the Wyant laboratory, Malhotra laboratory (MGH), R. Soberman (MGH), and R. Wolfson (HMS), for helpful discussions and critical reading of the manuscript. Special thanks to the Mass General Brigham Nikon Center of Excellence for helpful discussions and imaging experimental design. S.Q., together with G.A.W, performed and designed experiments, and analyzed and assembled data. C.L. performed imaging and mRNA analysis with the help of S.Q. M.S. performed mouse biochemical experiments with the help of S.Q. and G.A.W. G.A.W. wrote the article.

## Resource Availability

Additional information or requests for resources will be fulfilled by the lead contact, Gregory Wyant. All plasmids generated in this study are available via addgene or fulfilled by the lead contact. RNA-sequencing data will be deposited in the NCBI Gene Expression Omnibus (Geo) and will be publically available upon manuscript acceptance for publication. This paper does not report original code.

## Methods

### Cell Culture

C2C12 mouse myoblasts were originally obtained from the American Type Culture Collection (ATCC). 293FT human embryonic kidney cells were originally obtained from Invitrogen. 293FT and C2C12 cells were maintained in Dulbecco’s minimum essential medium (DMEM) supplemented with 10% fetal bovine serum (FBS), penicillin (100 U/ml), and streptomycin (100 ÿg/ml). All experiments using C2C12 cells were performed with passage 4-6 cells. Fresh aliquots of 293FT were thawed every 4 weeks. Virally infected cells were selected with puromycin (5 µg/ml) or blasticidin (10 µg/ml) as appropriate for the vector used. All mammalian cells were grown in a humidified atmosphere containing 21% oxygen and 5% CO_2_ at 37 °C unless otherwise stated.

### Chemicals

MitoTracker DeepRed FM (CST) was prepared at a 1 mM stock concentration in DMSO and was diluted in cell culture media at a final concentration of 500 nM. LysoTracker Blue DND-2 (Thermo) came prepared at a 1 mM stock concentration and was diluted in cell culture media at a final concentration of 100 nM. Water soluble dexamethasone (Sigma) was prepared in water at a final concentration of 3 mg/kg.

### Antibodies

Primary antibodies used were: Rabbit anti-p70 S6 Kinase (CST #9202), Mouse anti-b-Actin (Santa Cruz sc-47778), Rabbit anti-HIF1A (CST #14179), Mouse anti-LC3B (CST #83506), Mouse Anti-Keima-Red (MBL International, M182-3M), Rabbit anti-COPB1 (ProteinTech 27469-1-AP), Rabbit anti-PHD2/EGLN1 (CST #4835S), Rabbit anti-BNIP3 (CST #3769, mouse specific), Rabbit anti-S6 (CST #2217), Rabbit anti-Foxo3A (CST #48648), Rabbit anti-Foxo1 (CST #2880), Mouse anti-MuRF1 (Santa Cruz sc-398608), Mouse anti-MAFbx/Atrogin-1 (Santa Cruz sc-166806), Rabbit anti-ARNT (CST #5537), Rabbit anti-HA (CST #3724), Rabbit anti-FUNDC1 (CST #49240), All primary antibodies used for immunoblot were diluted at 1:500 in TBS-T+5% BSA.

### cDNA Synthesis and Quantitative PCR analysis (qPCR)

Total RNA was extracted from C2C12 cells and mouse tissues using TRIzol Reagent (Life Technologies, 15596). cDNA was generated by reverse transcription using High Capacity cDNA synthesis kit (Thermo) according to manufacturer’s instructions. qPCR was performed using a QuantStudio 6 (Thermo) using Sybrgreen or Taqman (Thermo) according to the manufacturer’s instructions. All quantitative calculations were performed using the 2^-DDCt^ method using *b-actin* or *Gapdh* as a reference gene.

### The following mouse Sybrgreen qPCR primers were used

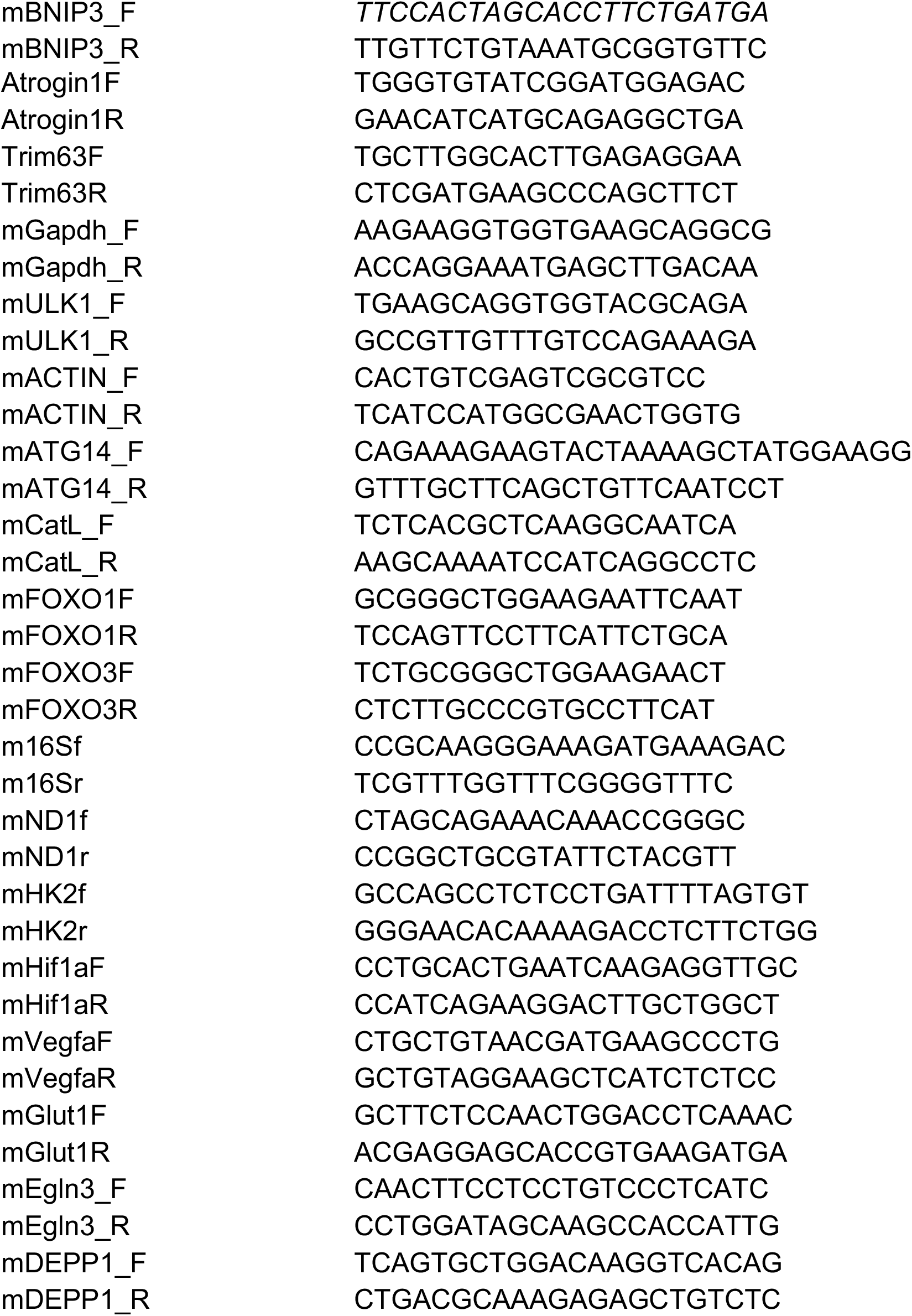

The following Taqman Probes were used:

Taqman Probes Used in this study

The following Taqman Probes were used: Taqman Probes Used in this study
| S.No | Catalog No. | Gene Name |
| --- | --- | --- |
| 1 | Mm00437306_m1 | Vegfa |
| 2 | Mm044225274_s1 | mt-ND1 |
| 3 | Mm04260181_s1 Rnr2 | mt-16s |
| 4 | Mm01185221_m1 | Trim63 |
| 5 | Mm00472200_m1 | Egln3 |
| 6 | Mm04206895-m1 | DEPP1 |
| 7 | Mm02619580_g1 Actb | b-Actin |
| 8 | Mm01275600_g1 BNIP3 | Bnip3 |
| 9 | Mm00468869_m1 Hif1a | Hif1a |
| 10 | Mm00441480_m1 Slc3a1 | Glut1 |
| 11 | Mm00443385_m1 HK2 | HK2 |
| 12 | Mm00499523_m1 Fbx032 | Atrogin-1 |

### RNA Sequencing

RNA was extracted from the gastrocnemius muscle 12-week-old mice of indicated genotypes. Each group comprised of 3-4 individual mice of which RNA was extracted using using TRIzol Reagent (Life Technologies, 15596). RNA was dissolved in RNase-free water and submitted to Novogene for eukaryotic mRNA-sequencing. RNAseq data was analyzed and p-values calculated using a negative binomial statistical model as implemented in DESeq.

### CRISPR/Cas9 Plasmid Generation

The LentiCRISPR_v2-puromycin or -blasticidin vectors were used to express sgRNAs with Cas9. LentiCRISPR_v2 was digested with BsmBI, gel purified and ligated with annealed oligonucleotides. Sense and antisense oligonucleotides corresponding to the desired sgRNA were mixed at equimolar ratios (0.25 nanomoles of each sense and antisense oligonucleotide) and annealed by heating to 100 °C in annealing buffer (1X T4 Ligase buffer, T4 PNK) followed by slow cooling from 95 °C to 4 °C. The annealed oligonucleotides were then diluted 1:200 in dH2O and ligated into the digested CRISPR vectors by incubation with T4 DNA ligase for 30 minutes at room temperature. The ligation reaction was transformed into XL-10 Gold ultracompetent cells and ampicillin-resistant colonies were verified by Sanger sequencing.

The following sgRNAs were used:

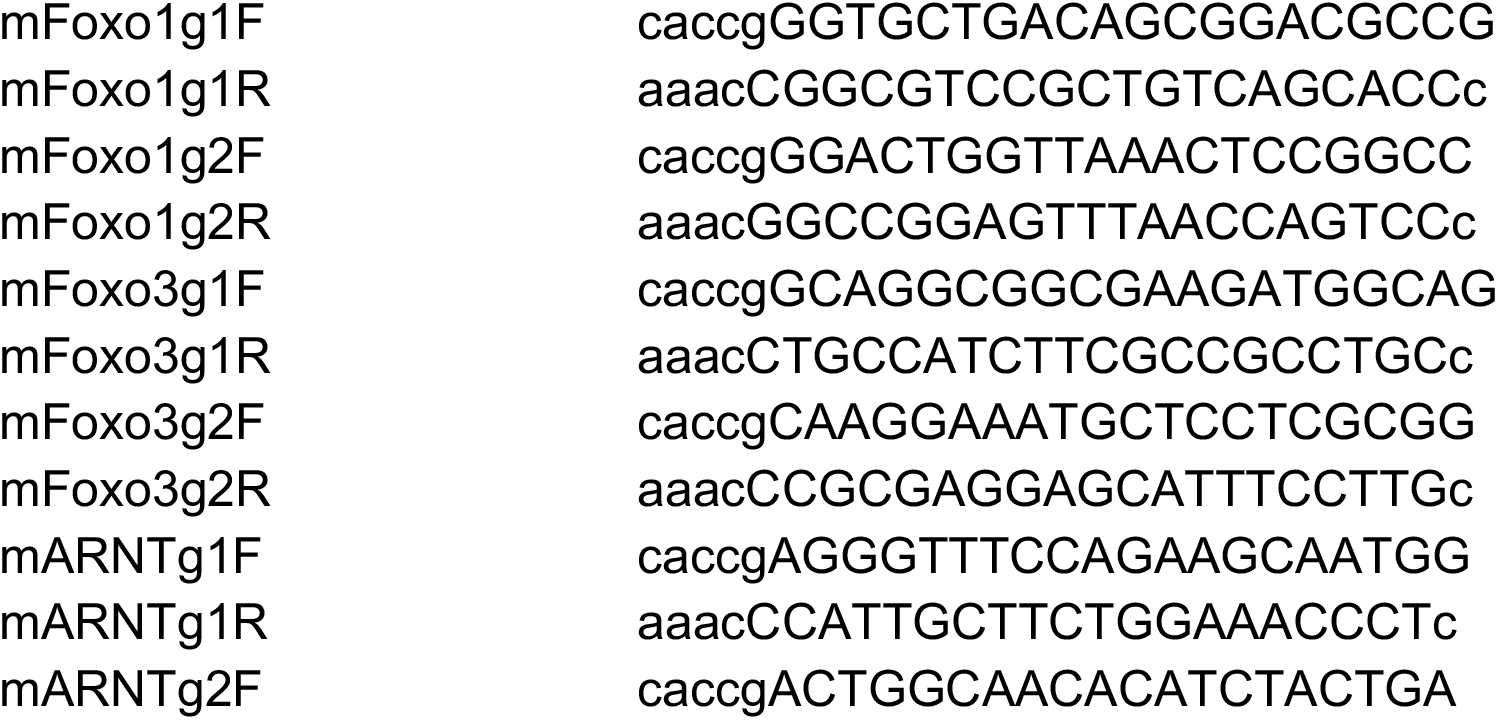
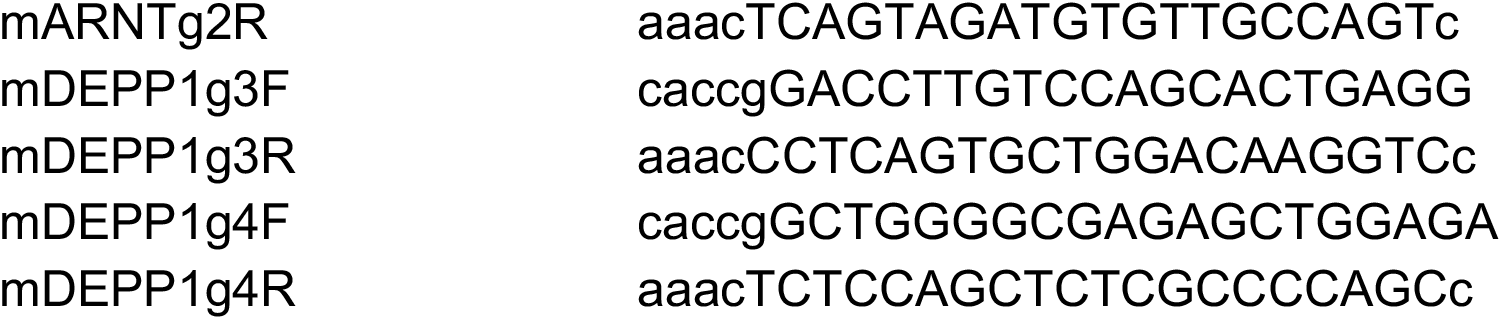

### Lentivirus Generation and Infection

To make the lentiviruses, 1.3 x 10^6^ HEK293FT cells were seeded in a 6-cm plate in DMEM supplemented with 10% FBS. Twenty-four hours later the cells were transfected with 1 ug of the desired lentivirus encoding plasmid together with of the packaging plasmids 0.5 μg psPAX2 and 0.5 μg pMD2.G using XtremeGene9 (Sigma) transfection reagent. Twelve hours after transfection, the medium was aspirated and replaced with 4 mL of fresh medium. Thirty-six hour later the virus-containing supernatants were collected and passed through a 0.45-μm filter to eliminate contaminating mammalian cells.

For lentiviral infections, 1 x 10^6^ target cells were combined with 250 μl of virus in 2 mL total volume of medium supplemented with 8 ug/mL polybrene and plated in 6-well plates. The plates were then immediately spun in an Eppendorf 581R centrifuge at 2,200 r.p.m. for 45 minutes at 37 °C. Twelve hours after infection, the virus-containing media was aspirated and replaced with fresh DMEM+10% FBS supplemented with Pen/Strep. Twenty-four hours post infection, the infected cells were trypsinized and replated in media that contained 5 ug/mL puromycin or 10 ug/mL blasticidin as appropriate for that virus.

### Immunoblot Analysis

Cells grown in 10-cm tissue culture dishes were washed once on ice with ice cold 1x phosphate-buffered saline (PBS) and then 1 mL ice cold PBS was added to each plate. The cells were then detached by scraping, transferred to a 1.5 mL Eppendorf tube, and pelleted at 1500 x g for 1 minute at 4 °C. The PBS was then aspirated, and the cell pellets were resuspended in lysis buffer (40 mM Hepes pH 7.4, 150 mM NaCl, 1.5 mM MgCl_2_, 1% Triton-X 100, Complete Mini EDTA-free Protease Inhibitor (Sigma), and Phosstop Phosphatase Inhibitor (Sigma)) and lysed by gentle rocking for 20 minutes at 4 ÿC. For samples involving mouse tissues, the entire mouse gastrocnemius, quadricep, or TA muscle were isolated and washed once in ice cold PBS. Following, the tissues were placed in an Eppendorf tube containing 1 mL RIPA buffer and homogenized using a Qiagen TissueLyser II for 10 minutes on the highest setting at 4 ÿC. For experiments involving Keima-fusion proteins, cell pellets were resuspended in RIPA buffer and lysed by gentle rocking for 20 minutes at 4 ÿC. The lysate was then clarified by centrifugation at 17,000 x g for 10 minutes at 4 ÿC and transferred to a new Eppendorf tube. The protein concentration of the whole cell or tissue extract was measured using the Bradford Assay and then normalized to 2 mg/mL. After normalization, the whole cell extract was denatured by the addition of 2.2% SDS, 11% glycerol, 100 mM DTT, and bromophenol blue.

Samples were resolved by SDS-polyacrylamide gel electrophoresis using 4-20% or 4-12% Tris-Glycine gels (Novex) and transferred onto nitrocellulose membranes using a Bio-Rad Trans-Blot Turbo Transfer system using the high molecular weight setting. Following transfer, membranes were incubated with Ponceau S staining solution (CST) for 3 minutes rocking at room temperature to visualize transferred bands following by washing in Tris-buffered saline +0.1% Tween 20 (TBS-T). Membranes were then blocked by incubation in 5% milk/tris-buffered saline + 0.1% Tween 20 with gentle rocking for 1 hour at room temperature. The membranes were washed with TBS-T prior to overnight incubation with primary antibody diluted in TBS-T+5% BSA with gentle rocking at 4 ÿC. After primary antibody incubation, membranes were washed three times with TBS-T and then incubated with horseradish peroxidase (HRP)-conjugated secondary antibody (CST) (1:5000) in 5% milk/TBS-T with gentle rocking for 1 hour at room temperature. The membranes were washed three times with TBS-T and bound antibodies were then detected with enhanced chemiluminescence western blotting reagents (Thermo Fisher Scientific, no. WBKLS0500) or Super-Signal West Pico (Thermo Fisher Scientific, no. PI34078).

Quantification of immunoblots was performed in ImageJ using the analyze gel function and are shown as relative values compared to the control or untreated group. For mouse experiments, each band was quantified individually. The band intensities for the control group were averaged, and each mouse was then normalized to the average band intensity of the control group. Band intensity for each mouse was then normalized to the control protein b-actin or S6. Values are shown as fold change versus control.

Quantification of LC3B immunoblots is shown as the ratio of LC3B-II: LC3B-I. Both bands of LC3B were quantified individually and each ratio was normalized to the wild-type ad libitum fed normal atmospheric oxygen control group.

For quantification of Keima processing experiments, the ratio of “processed Keima: Full length Keima fusion protein” is shown. Each ratio was normalized to wild-type cells grown in full media at normoxia. Both bands were quantified individually.

### Mice

The *Atg7^fl^*^/fl^ (JAX stock #034429), Egln1*^fl^*^/fl^(JAX stock #009672), *CAG-RFP-EGFP-LC3* (JAX stock #027139), Rosa26 HIF2 DPA (JAX stock #009674), and HSA-Cre79 (Acta-Cre, JAX stock #006149) mice were imported from Jackson Laboratory and were previously described. Experimental protocols were approved by the Mass General Brigham Institutional Animal Care and Use Committee. Animals were housed at 24°C in a 12-hour light/12-hour dark cycle where food and water were accessible ad libitum. All mouse experiments were performed on littermate male mice aged 6-15 weeks. Mouse experiments involving Egln1*^fl^*^/fl^, *Atg7^fl^*^/fl^, and Rosa26 HIF2 DPA mice utilized Cre expressing wild-type mice to control for possible Cre toxicity.

To generated Depp1-Tg mice, a targeting vector was generated via Ingenious Targeting Laboratories. The Depp1 cDNA sequence was fused to an in-frame HA tag and cloned using restriction digest methods into the vector backbone pROSA26-pCAG. The Depp1-HA sequence is driven by a pCAG promoter upon Cre excision of the LoxP Neo-Stop cassette. The targeting vector contains a short homology arm of 1.1 kilobases (kb) of the ROSA26 genomic sequence and a 4.3 kb long homology arm of downstream ROSA26 sequence of a total vector size of ∼18 kb. The resulting vector was then linearized using PvuI prior to electroporation into C57BL/6 ES cells using the Harvard Medical School Transgenic Mouse Core. Once positive clones were identified, they were subsequently injected into blastocysts to generate male chimeras. Heterozygous mice were obtained by breeding the chimeras with wild-type C57BL/6 females and the resulting pups were backcrossed with wild-type mice for 10 generations.

Depp1^-/-^mice were described previously and generated by utilizing two guide RNAs (sgRNAs) targeting the mouse Depp1 gene using CRISPR-Cas9 injection into C57/BL6 mouse zygotes. In brief, to generate mDEPP1 sgRNA, a gblock with a built-in T7 priming sequence and the guide/scaffold sequence was synthesized from IDT using the following sequence:

mDEPP1g1 T7:

cgctgTTAATACGACTCACTATAGGGCCTCAGTGCTGGACAAGGTCGTTTTAGAGCTAGAAAtagcaagttaaaataaggctagtccgttatcaacttgaaaaagtggcaccgagtcggtgcTTTT

mDEPP1g2 T7:

cgctgTTAATACGACTCACTATAGGGGGAGAGCAGGCAGATGGGAGGTTTTAGAGCTAGAAAtagcaagttaaaataaggctagtccgttatcaacttgaaaaagtggcaccgagtcggtgcTTTT

Depp1-HA mice were generated using a single guide RNA targeting the 3’end of the mouse Depp1 gene and an HDR repair oligo containing an in-frame linker and HA tag sequence. The sgRNA PAM site within the HDR repair oligo was mutated to a silent, synonymous codon. To generate the Depp1-HA sgRNA, a gblock with a built-in T7 priming sequence and the guide/scaffold sequence was synthesized from IDT using the following sequence:

mDepp1HA:

cgctgTTAATACGACTCACTATAGGGAGAGTTCATGGATCACTGGGGTTTTAGAGCTAGAAAtagcaagttaaaataaggctagtccgttatcaacttgaaaaagtggcaccgagtcggtgcTTTT

The following HDR oligo was used: GAAAAGCTGGACTTCTAGAAAGTCCTGCCGGGCCTTAGCGTCCGTCTCCAGCTCTCGCCCCAGCAGTATCCTAGGTACTCTCTATTTGCATCTCCCAGTGATCCATGAACTCggagggagcggaGCTAGCTACCCATACGATGTTCCAGATTACGCTTAACTCCTACCCCAGGAAAAACCCATAAAGGATATGGTGGGCC

The lyophilized gblock was reconstituted in 10 mL PCR-grade water to make a 20 ng/mL stock. RNA was synthesized in vitro from the gblock using MEGAshortscript T7 Kit (LifeTech) using 8 mL of the reconstituted gblock as a template. To capture the RNA, 50 mL Agencourt RNAClean XP beads (A63987) were added to the synthesized RNA and incubated at room temperature for 10 minutes. The beads were then collected using 96-well plate magnet and washed 3 times in 80% EtOH. After the final EtOH wash, the ethanol was aspirated, and the beads were left to dry and resuspended in HyClone Molecular Biology-Grade Water (GE Healthcare Life Sciences). The beads were then placed back on the magnet and the supernatant was collected. The RNA concentration was quantified by nanodrop and then immediately frozen at -80 ÿC until injection.

Zygote injections were performed at the HMS Transgenic Facility. All procedures were performed according to National Institutes of Health guidelines and approved by the Committee on Animal Care at HMS. Female 8–10-week-old C57/BL6 mice were superovulated by IP injection of 5 IU of pregnant mare serum gonadotropin (367222-1000IU; EMD Millipore) followed 46–48 hours later by 5 IU human chorionic gonadotropin (80051–032; VWR). Superovulated female mice were then mated to stud males. Fertilized pronuclear stage embryos (zygotes) were collected ∼20 hours after injection of human chorionic gonadotropin. Cytoplasmic injections were performed using a Piezo actuator (PMM-150FU; Prime Tech) and a flat-tip microinjection pipette with an internal diameter of 8 µm (Origio). The injection mix was prepared immediately before the procedure and included the following components at the final concentrations indicated: 100 ng/µl Cas9 mRNA (CAS9MRNA-1EA; Sigma Aldrich), and 50 ng/µl sgRNA. Immediately after the completion of the injection, zygotes were transferred into the oviducts of pseudopregnant females at 0.5 dpc.

For genotyping *Depp1*^-/-^ animals, tails snips were collected into tubes with 500 µl QuickExtract buffer (Epicentre). To obtain PCR-ready genomic DNA, the tubes were incubated at 65°C for 10 minutes, followed by a quick vortex and a 2-minute incubation at 98 °C. The insoluble material was removed by centrifugation at 13,000 x g for 10 minutes at 4 ÿC. To genotype by PCR, 5 µl of supernatant was used as a PCR template and the following primers were used

mDEPP1Fwd: 5’-tcgccactggtttcctcttg -3’

mDEPP1Rev: 5’-GAGTTCATGGATCACTGGGAGG-3’

For genotyping the Depp1-HA animals, the following primers were used

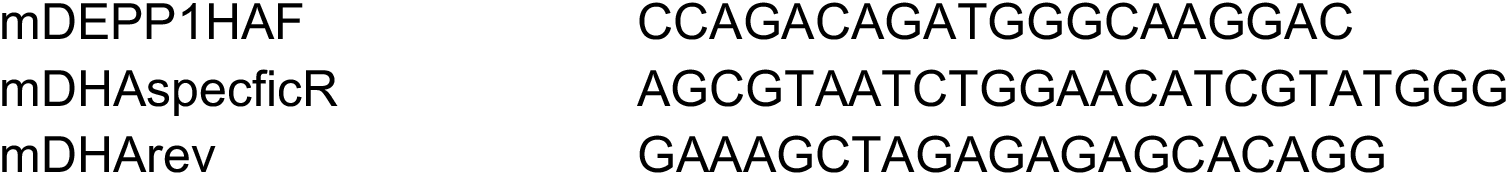

PCR was done with Q5 High Fidelity 2X Master Mix (New England Biolabs) at 98 °C for 30 seconds, 30 cycles of 98 °C for 10 seconds, 64 °C for 30 seconds, and 72 °C for 30 seconds, and a final extension time of 72 °C for 2 minutes. PCR products were sequenced by Amplicon sequencing.

### Fasting-induced muscle atrophy

For experiments involving mouse fasting, mice to be fasted were moved into a clean cage with new bedding with free access to water and deprived of food for 48 hours prior to euthanasia.

### Hypoxia-induced muscle atrophy

For experiments involving hypoxia, mice housed at hypoxia were placed in a hypoxia chamber at 10% O_2_ for 7-days or indicated time and food, water, and bedding was refreshed every other day prior to euthanasia.

### Dexamethasone-induced muscle atrophy

For experiments involving dexamethasone, water soluble dexamethasone (3mg/kg) was added to the drinking water of the mouse cage for a period of 14-days.

### Immunohistochemistry and tissue immunofluorescence

All tissues were fixed with buffered 10% formalin solution (SF93-20; Fisher) followed by 70% Ethanol. For hematoxylin and eosin, tissues were embedded in paraffin prior to sectioning.

For fluorescent microscopy imaging on tissues, freshly harvested tissue samples were fixed in 4% paraformaldehyde for 12 hours, washed 3 times with 1x PBS and cryoprotected in 30% sucrose/1x PBS overnight at 4°C before embedding in O.C.T. (Tissue-Tek^®^, Sakura Finetek, Cat# 4583) and snap freezing in isopentane.

### Confocal Microscopy

Live cell confocal microscopy were performed using a Zeiss LSM 900 confocal microscope equipped using a 63 x Plan-Apochromat oil objective, NA 1.4. For live cell imaging experiments, 2 x 10^5^ cells were plated onto 35 mm-glass bottom dishes (No. 1.5, 14 mm glass diameter, MatTek) then incubated in phenol-red free medium (FluoroBriteDMEM, Thermo Fisher) containing 10% fetal bovine serum. At time of imaging, cells were imaged in FluoroBrite DMEM without 10% fetal bovine serum. Series optical sections were collected with a step-size of 0.22 microns. Z series were displayed as maximum z-projections, and gamma, brightness, and contrast were adjusted for each image equally using FiJi software. At least 10 image frames per condition were analyzed without exclusion.

### Seahorse analysis of oxygen consumption rate (OCR)

C2C12 cells (10,000 cells/well) were plated in XF-96 plates. The next morning, the cells were washed once with 1xPBS and incubated overnight in serum-free DMEM. The cells were then washed once and incubated for 1 hour in XF assay medium (DMEM pH 7.4 with 10 mM glucose, 2 mM L-glutamine, 1 mM pyruvate) in a non-CO_2_ incubator per manufacturer’s instructions (Seahorse Agilent). Real-time measurements of extracellular acidification rate (ECAR) and oxygen consumption rate (OCR) were performed using an XF-96 Extracellular Flux Analyzer (Agilent). Three or more consecutive measurements were obtained under basal conditions and after the sequential addition of 1.5 mM oligomycin to inhibit mitochondrial ATP synthase; 1 mM FCCP (fluoro-carbonyl cyanide phenylhydrazone), a protonophore that uncouples ATP synthesis from oxygen consumption by the electron-transport chain; and 500 nM rotenone plus antimycin A, which inhibits the electron transport chain.

### Statistical Analysis

GraphPad Prism version 10 was used for statistical analysis included in the main and supplementary figures. For all experiments, statistical significance was calculated using unpaired, two-tailed Student’s *t* test when comparing two groups or one-way ANOVA when comparing three or more groups. *P* values were considered statistically significant if the *P* value was <0.05. For all figures, * indicates *p* < 0.05 unless otherwise indicated. Error bars represent SD unless otherwise indicated. Data was evaluated for normality using the Shaprio-Wilk test and D’Agostino & Pearson Test. No statistical methods were used to pre-determine the sample size. No sample size calculation was performed for fluorescence microscopy experiments. All RNA-sequencing measurements were performed on 3-4 independent biological replicates for each condition.

## Supplementary Figure Legends

**Supplementary Figure 1:**
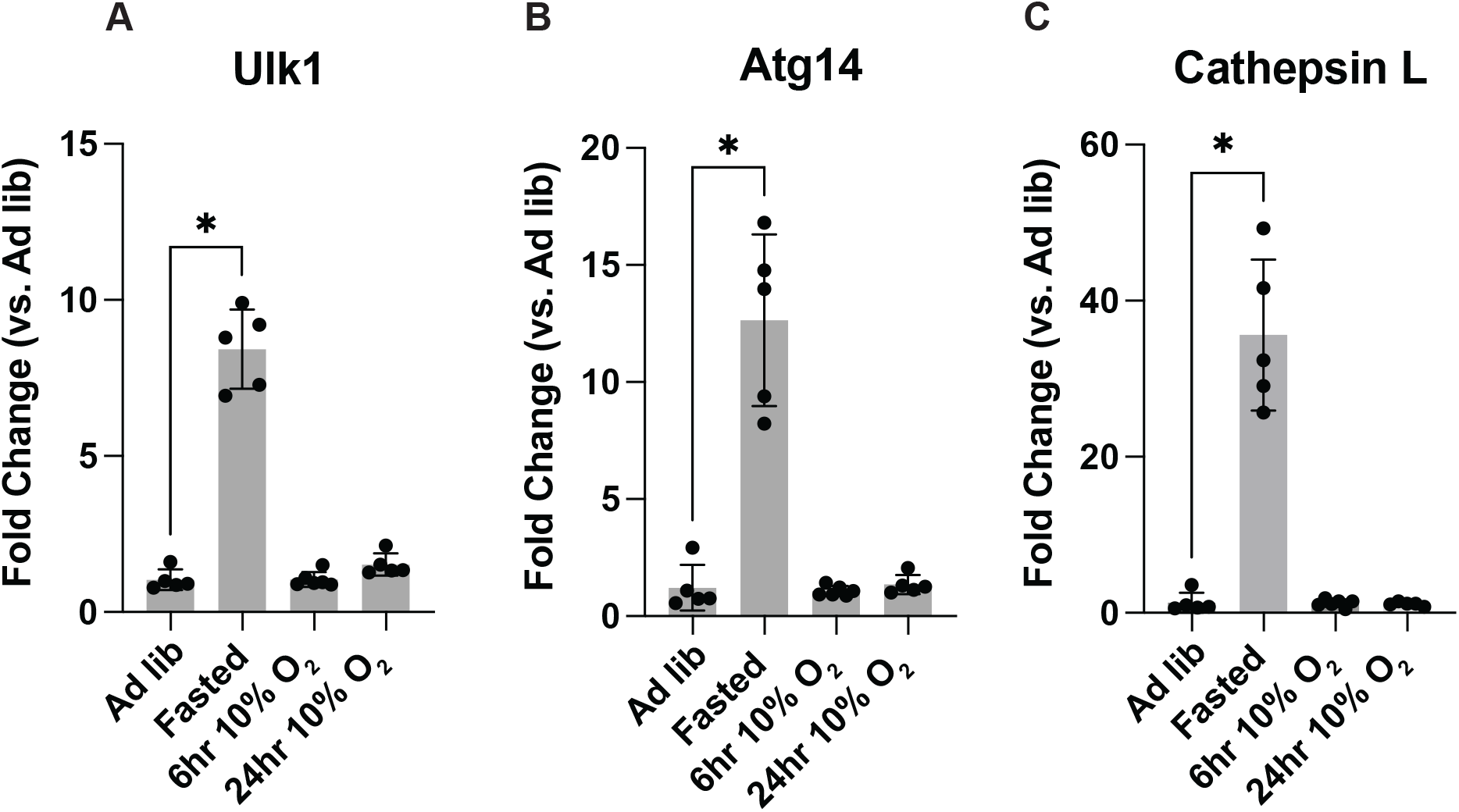
Hypoxia does not increase multiple known atrogenes in skeletal muscle. **(A-C)** mRNA analysis in gastrocnemius muscles isolated from mice deprived of food for 48 hours (starvation), housed at 10% O_2_ (hypoxia) for 6 hours or 24 hours, or fed ad libitum and housed at normal atmospheric oxygen (21% O_2_). * indicates P<0.05. N=3 per group.

**Supplementary Figure 2:**
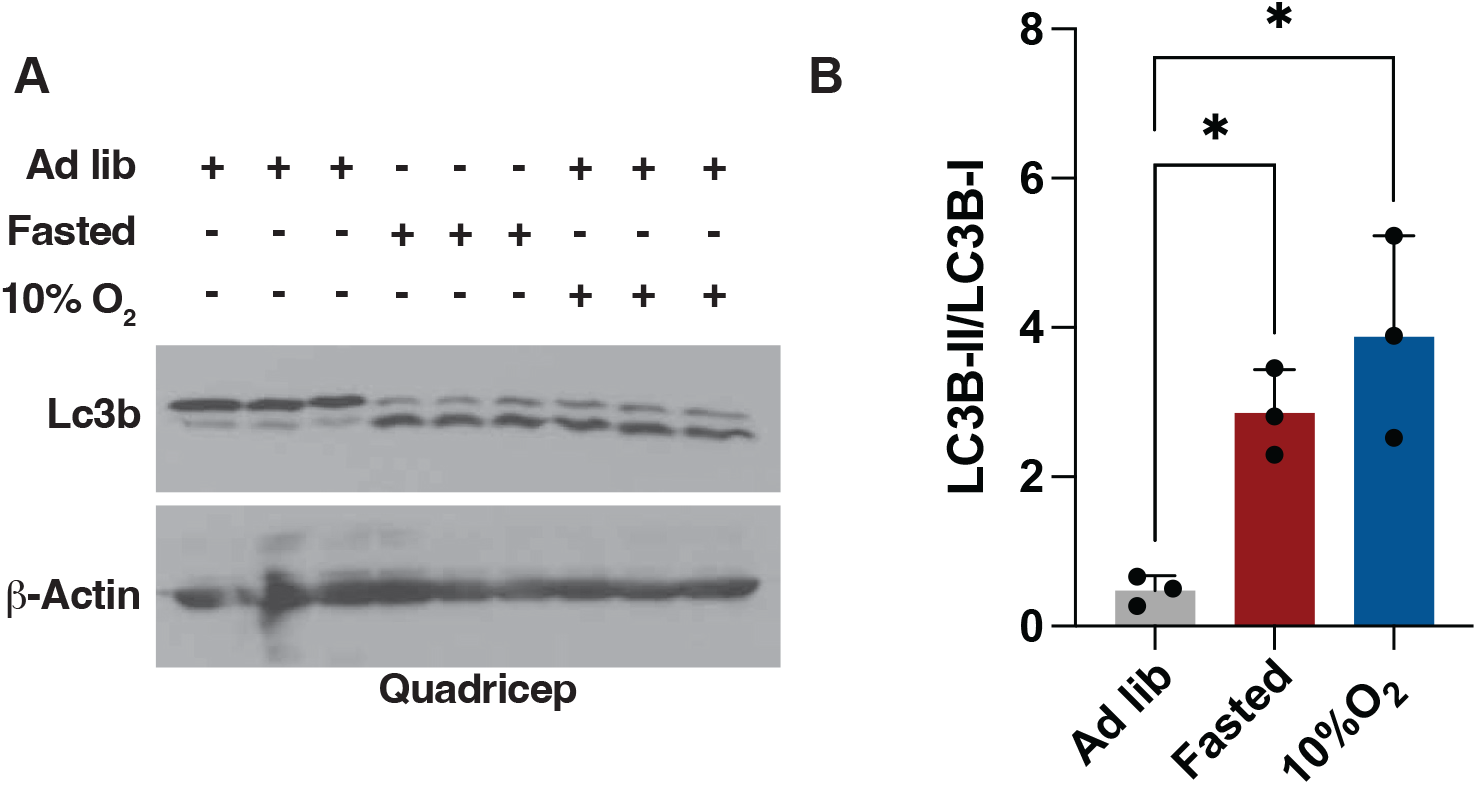
Fasting or hypoxia promotes Lc3b lipidation in skeletal muscle. **(A-B)** Immunoblot (A) and densitometry (B) analysis of mouse quadricep muscle lysates isolated from mice either deprived of food (fasted) for 48 hours, housed at 10% O_2_ for 7 days, or provided food ad libitum and housed at normal atmospheric oxygen tension (adlib) prior to euthanasia. Each sample represents individual mouse. * indicates P<0.05, N=3.

**Supplementary Figure 3:**
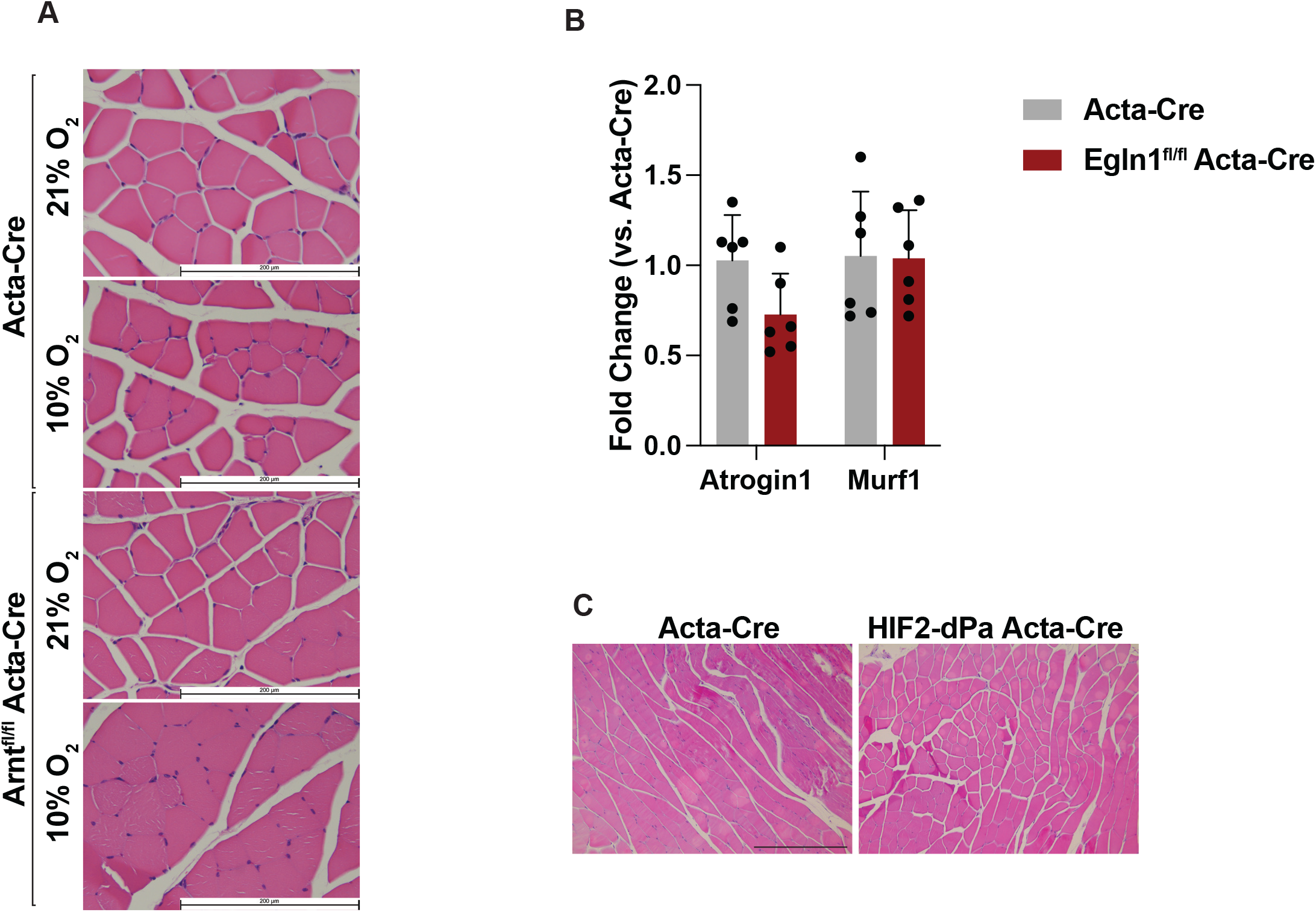
HIF and muscle atrophy. **(A)** Hematoxylin and Eosin (HE) staining of gastrocnemius muscle sections isolated from Acta-Cre (Wild-type) and Arnt^fl/fl^ Acta-Cre mice. Where indicated, mice were housed at 10% O_2_ or normal atmospheric oxygen (21% O_2_) for 7 days prior to euthanasia. **(B)** mRNA analysis in gastrocnemius muscles isolated from Acta-Cre (Wild-type) and Egln1^fl/fl^ Acta-Cre mice. N>3 per group. **(C)** Hematoxylin and Eosin (HE) staining of gastrocnemius muscle sections isolated from Acta-Cre (Wild-type) and HIF2-dPA Acta-Cre mice.

**Supplementary Figure 4:**
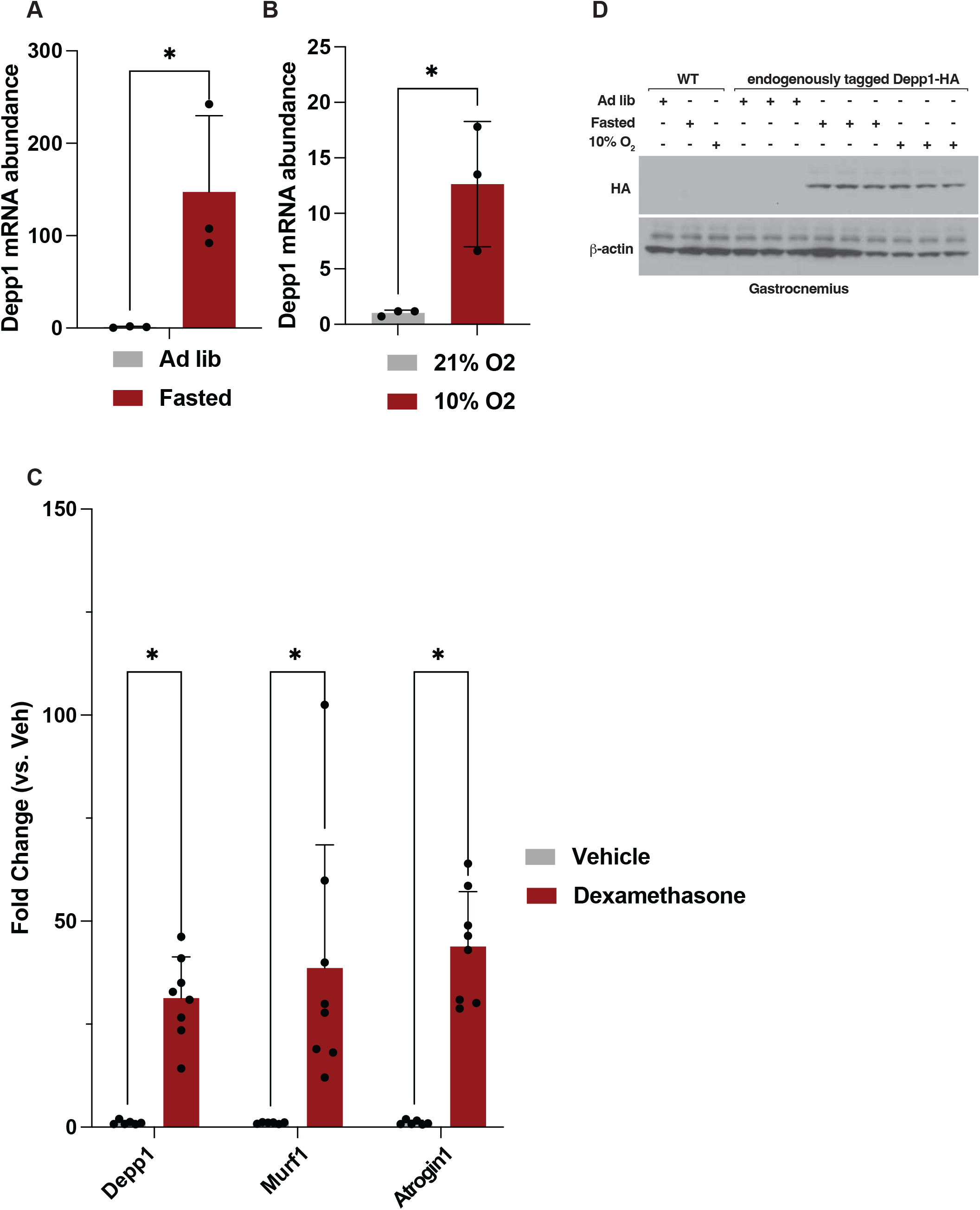
Depp1 regulation in vivo. **(A-B)** Depp1 mRNA analysis in gastrocnemius muscles isolated from mice fasted for 48 hours (A), housed under 10% O_2_ for 18 hours (B), or fed ad libitum and housed at normal atmospheric oxygen (21% O_2_). * indicated P<0.05. N=3. **(C)** mRNA analysis in gastrocnemius muscles isolated from mice treated with dexamethasone (3 mg/kg) or vehicle for 14 days prior to RNA extraction. * indicates P<0.05. N>3 per group. **(D)** Immunoblot analysis of mouse gastrocnemius muscles isolated from wild-type or endogenously tagged Depp1-HA mice. Where indicated, mice were fasted for 48 hours, housed under 10% O_2_ for 18 hours (B), or fed ad libitum and housed at normal atmospheric oxygen (21% O_2_).

**Supplementary Figure 5:**
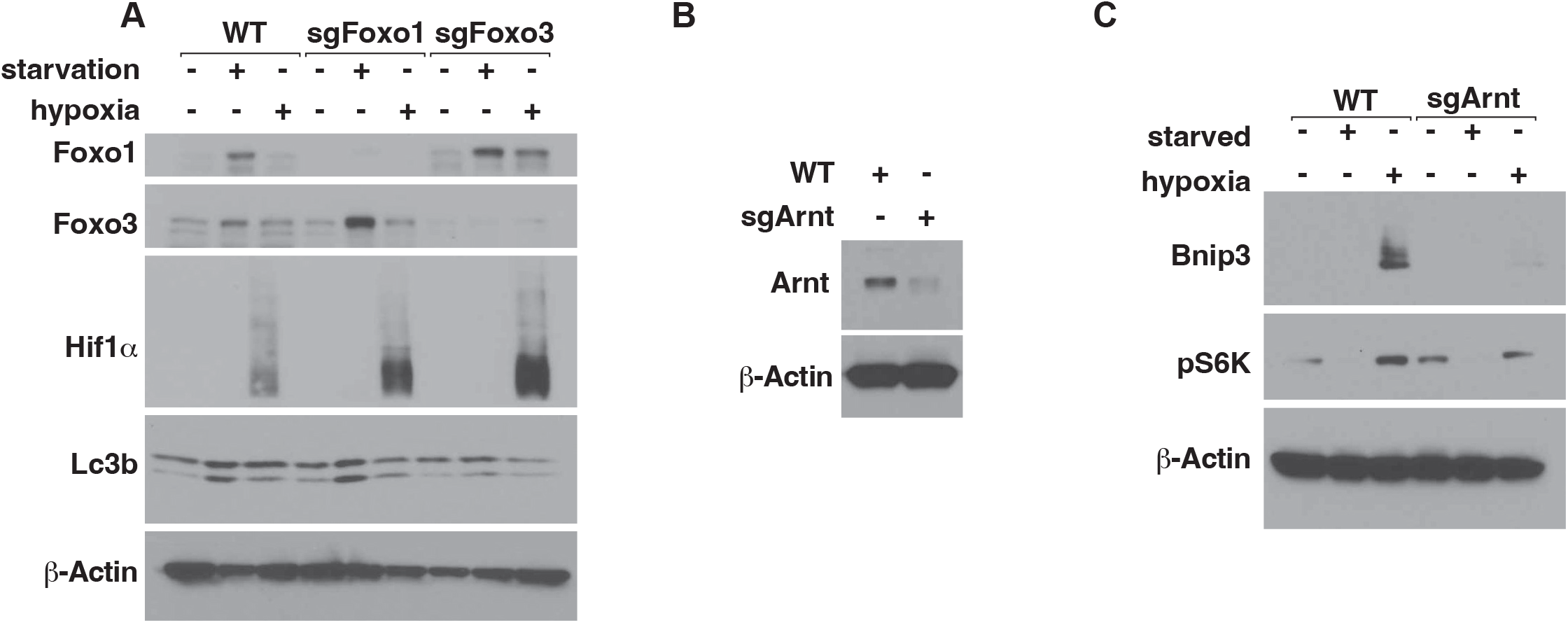
Validation of sgFoxo1, sgFoxo3, and sgArnt C2C12 cells. **(A-C)** Immunoblot analysis of wild-type (WT), sgFoxo1, sgFoxo3 (A), and sgArnt (B-C) C2C12 cells. Where indicated, cells were starved of nutrients, grown at 0.1% O_2_, or grown in full media at normal atmospheric oxygen tension for 18 hours prior to cell lysis.

**Supplementary Figure 6:**
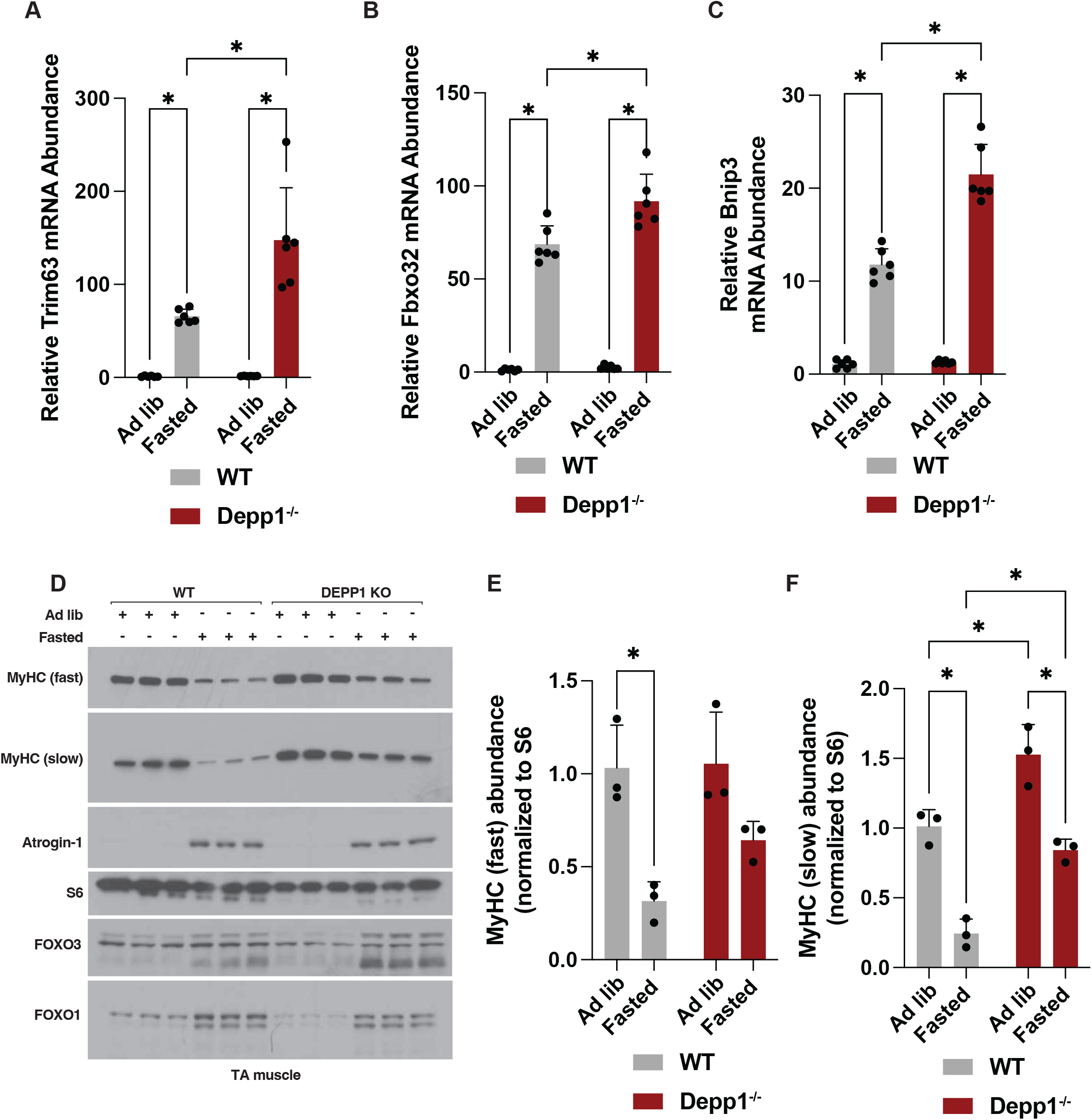
Mice lacking Depp1 spare Myosin Heavy Chain abundance upon fasting. **(A-C)** mRNA analysis in gastrocnemius muscles isolated from wild-type or Depp1^-/-^ mice fasted for 48 hours or fed ad libitum. * indicates P<0.05. N>3. **(D-F)** Immunoblot (D) and densitometry (E-F) analysis of tibialis anterior (TA) muscle lysates isolated from wild-type (WT) and Depp1^-/-^ mice. Where indicated, mice were fasted for 48 hours or provided food ad libitum prior to tissue lysis. * indicates P<0.05. N=3 per group.

**Supplementary Figure 7:**
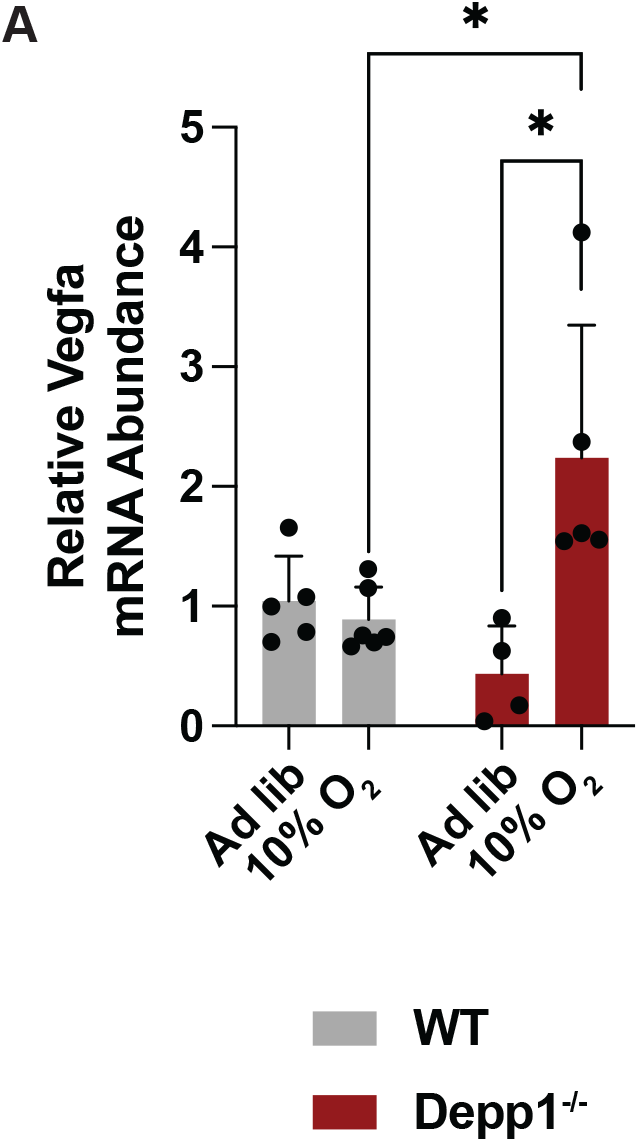
Increased Vegfa gene expression in Depp1^-/-^ mice. **(A)** mRNA analysis in gastrocnemius muscles isolated from mice housed under 10% O_2_ for 6 hours or housed at normal atmospheric oxygen (21% O_2_). * indicated P<0.05. N>3.

**Supplementary Figure 8:**
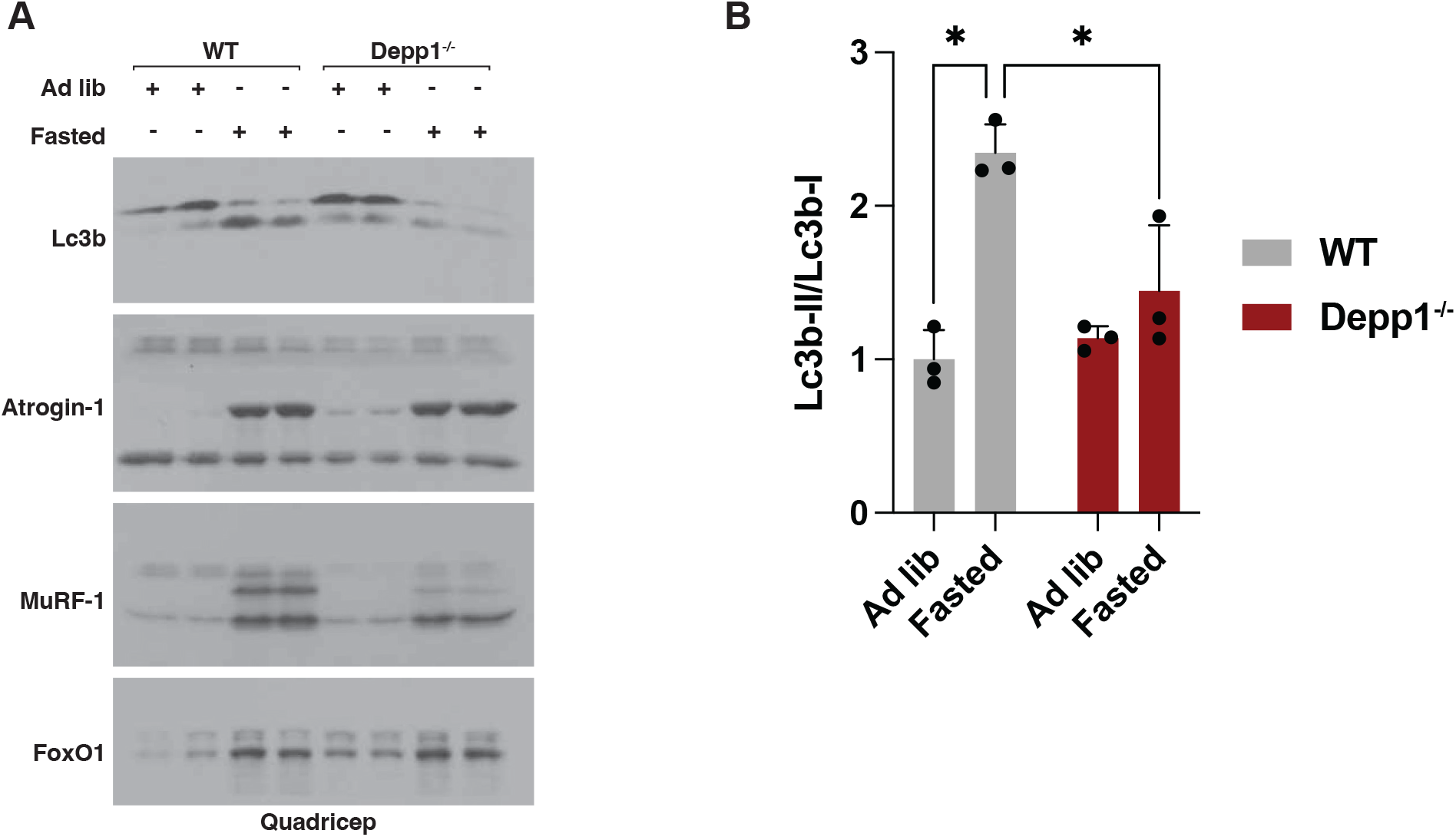
Depp1 loss reduces Lc3b lipidation under fasting in vivo. **(A-B)** Immunoblot (A) and densitometry (B) analysis of quadricep muscles isolated from wild-type and Depp1^-/-^ mice. Where indicated, mice were fasted for 48 hours or fed ad libitum. * indicates P<0.05. N=3.

**Supplementary Figure 9:**
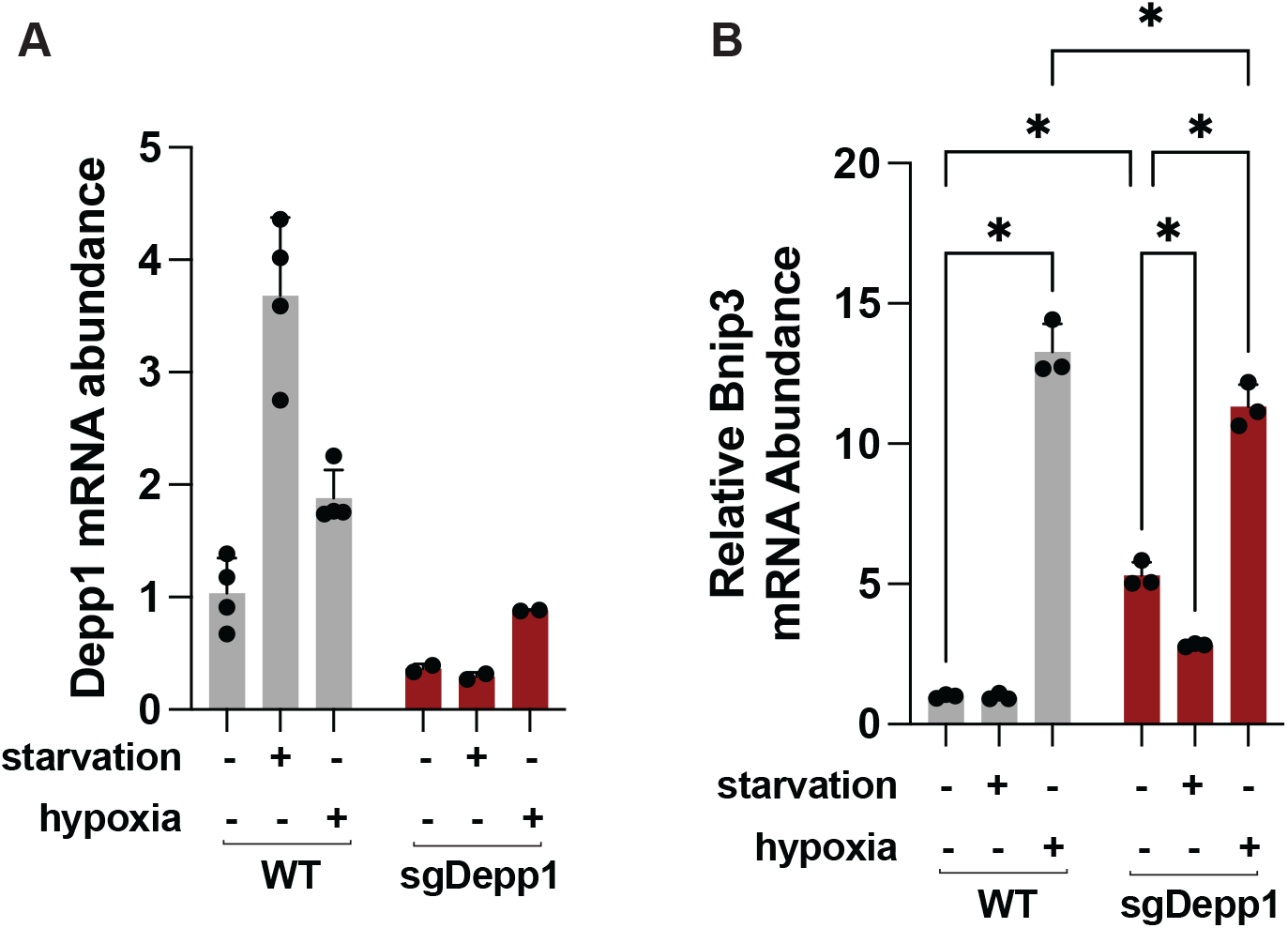
Validation of Depp1^-/-^ C2C12 cells. **(A-B)** mRNA analysis in wild-type and sgDepp1 C2C12 cells. Where indicated, cells were deprived of nutrients (starvation), grown at 0.1% O_2_ (hypoxia), or grown in full media at normal atmospheric oxygen tension for 18 hours prior to RNA extraction. * indicates P<0.05. In (A), N>2 per group while in (B) N=3 per group.

**Supplementary Figure 10:**
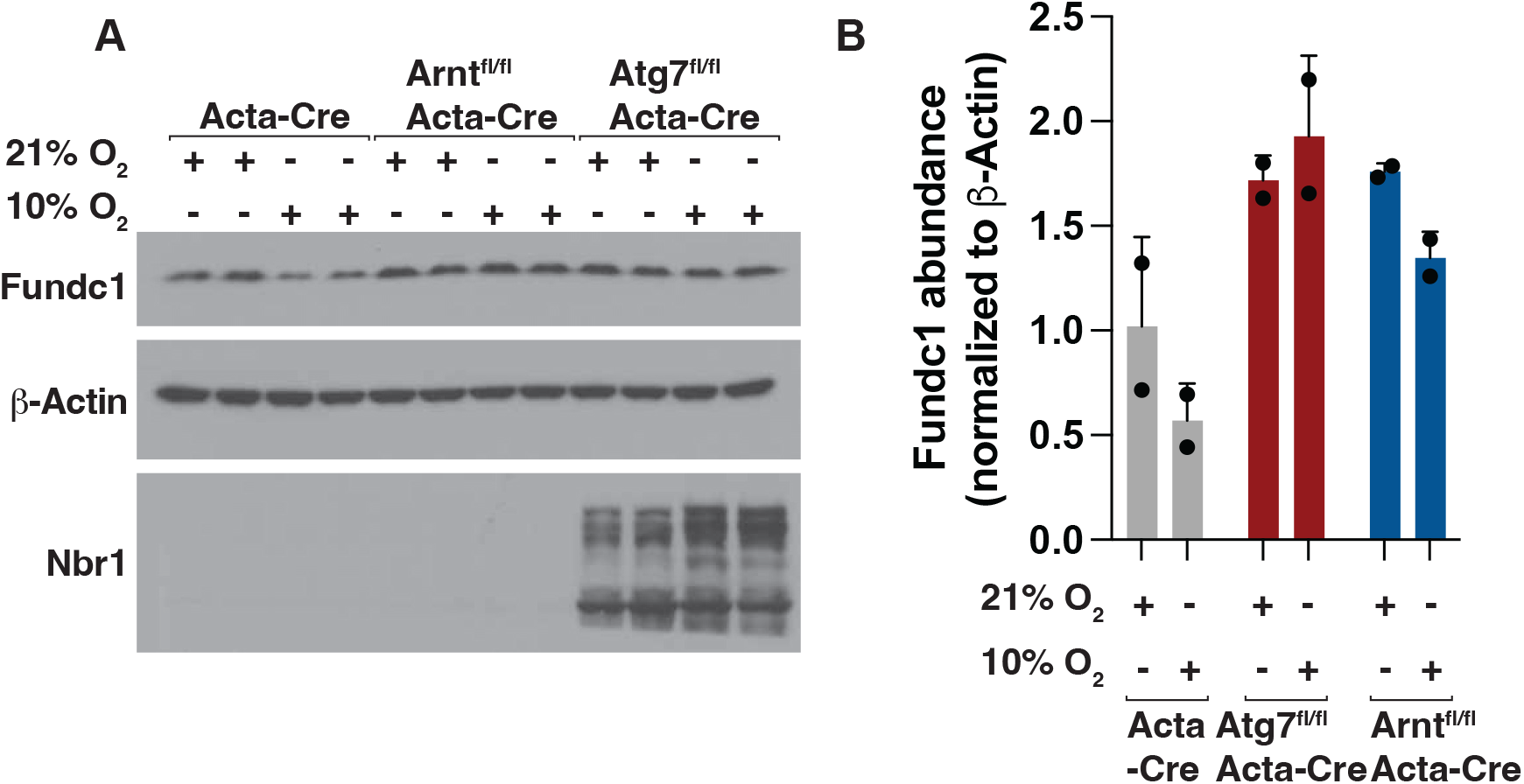
Hypoxia reduces Fundc1 abundance in a Hif and autophagy-dependent manner in vivo. **(A-B)** Immunoblot (A) and densitometry (B) analysis in quadricep muscle lysates isolated from Acta-Cre, Arntf^l/fl^-Acta-Cre, and Atg7^fl/fl^-Acta-Cre mice. Where indicated, mice were housed at 10% O_2_ (hypoxia) or 21% O_2_ (normal atmospheric oxygen tension) for 7 days prior to tissue lysis. N=2 per group.

